# Microbiome-Derived RNA Promotes Heritable Defense Against Intracellular Pathogens in *Caenorhabditis elegans*

**DOI:** 10.64898/2026.06.09.731142

**Authors:** Jordan D. West, Samuel A. Schwartz, Samuel Li, Peri Roper, Vladimir Lažetić

## Abstract

Intestinal microbiota shape host immunity, but whether they can pre-activate defense against intracellular pathogens remains unclear. Here, we identify a natural microbiome member, *Stenotrophomonas indicatrix* JUb19, as the first example of a bacterium that partially induces the Intracellular Pathogen Response in *Caenorhabditis elegans* in the absence of infection. This response is coupled to broader metabolic remodeling, suggesting that combined immune and metabolic changes contribute to defense. We show that bacterial single-stranded RNA contributes to this effect, identifying a novel microbiome-derived signal that links microbial exposure to host defense. JUb19 exposure reduces susceptibility to both viral and microsporidian pathogens and confers inherited protection to unexposed progeny, but at a cost to parental fitness. These findings reveal a mechanism by which microbiome-derived RNA programs immunity against intracellular pathogens across generations.

## Introduction

Intestinal microbiome composition influences infection outcome, yet how microbiota shape host defense programs, particularly against intracellular pathogens, remains poorly defined. The nematode *Caenorhabditis elegans* provides a powerful genetically tractable system for studying host–microbe interactions^1–3^. In nature, *C. elegans* associates with diverse bacterial communities that influence development, metabolism, and pathogen susceptibility, whereas under laboratory conditions, bacterial exposure can be precisely controlled to test the effects of defined microbes on host physiology^1,2,4–7^. Previous work has shown that natural bacterial isolates can alter susceptibility to extracellular intestinal pathogens through antimicrobial metabolites and modulation of host immune signaling^6–8^.

Intracellular infection poses distinct challenges because pathogens replicate within host cells, where many extracellular defenses are ineffective^9,10^. Natural microbiome variation has been associated with altered outcomes of intracellular infection, but these studies primarily measured pathogen burden rather than underlying immune-state changes^2,3^. In *C. elegans*, Orsay virus and the microsporidian pathogen *Nematocida parisii*, an obligate intracellular fungal-like parasite, trigger the Intracellular Pathogen Response (IPR), a transcriptional program characterized by strong induction of approximately 80 genes together with broader, lower-level transcriptional changes^11–15^. Although the molecular effectors of the IPR remain incompletely understood, activation of this program is associated with resistance to intracellular infection and proteotoxic stress^14,16–19^. The IPR provides a framework for studying epithelial cell immunity in an organism lacking professional immune cells and shares regulatory similarities with the type I interferon response in mammals^20,21^. The IPR is distinct from antibacterial immune programs and is not activated by extracellular bacterial pathogens^13,14^. Whether commensal bacteria can engage this intracellular defense pathway had not been previously explored.

In this study, using a defined collection of natural microbiome isolates^1^, we identified *Stenotrophomonas indicatrix* strain JUb19 as a bacterial species that induced expression of an IPR reporter and enhanced resistance to both Orsay virus and *N. parisii*. This protection extends to progeny, indicating that microbiome-derived signals can generate inherited defensive states. Rather than behaving as an intracellular pathogen, JUb19 activated host defense before substantial intestinal colonization, and bacterial RNA was sufficient to provide protection. Transcriptomic analyses revealed that JUb19 did not broadly activate the canonical IPR but instead induced a distinct immune–metabolic transcriptional state that combines selective signatures of the IPR with metabolic reprogramming. This protective state was accompanied by reduced fitness, suggesting a tradeoff between immune readiness and other physiological functions.

## Results

### *Stenotrophomonas indicatrix* (JUb19) and related *Stenotrophomonas* species induce IPR reporter expression

We screened 12 representative species naturally associated with *C. elegans* for their ability to induce expression of the IPR transcriptional reporter *pals-5*p::GFP (*jyIs8*). We found that *S. indicatrix* (strain JUb19) was the only species that induced GFP expression when fed to animals, compared to the standard laboratory diet of *Escherichia coli* (strain OP50).

Animals were first examined 48 hours (h) post-exposure at 20 °C, when both JUb19– and OP50-treated populations reached the L4 stage. IPR reporter expression in JUb19-fed animals was significantly higher than that in OP50-fed controls (Fig. 1a, b, Supplementary Table 1). To test whether this increase in GFP expression reflects reduced transgene silencing rather than bona fide IPR activation, we also measured the expression of the pharyngeal co-injection marker *myo-2*p::mCherry, whose expression is unrelated to IPR activation (Fig. 1c, Supplementary Table 1). We observed similar mCherry expression levels between treatments, suggesting that the GFP increase is IPR specific. We next examined animals at 72 h post-exposure, when they had reached adulthood. JUb19-treated animals again showed increased GFP expression, consistent with results from the L4 stage (Fig. 1d, e, Supplementary Table 1). However, we also observed a relatively small but significant increase in mCherry expression at this time point (Fig. 1f, Supplementary Table 1). Together, these results suggest that JUb19 is the first bacterial species capable of inducing IPR reporter expression in *C. elegans*.

**Figure 1:**
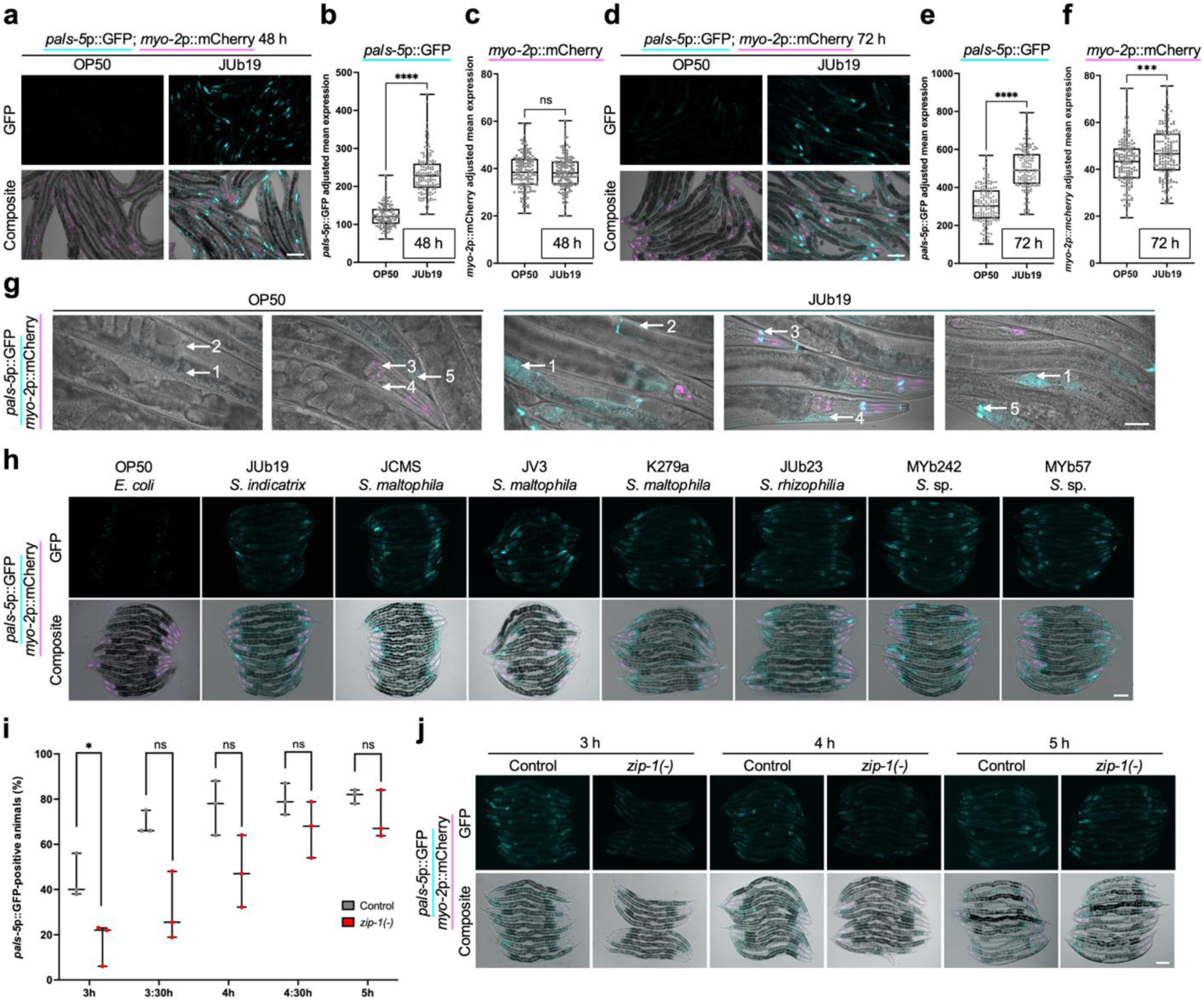
*S. indicatrix* (JUb19) and related *Stenotrophomonas* species induce IPR reporter expression. **a** Representative images of animals fed OP50 and JUb19 at 48 h. **b, c** Quantification of *pals-5*p::GFP (**b**) and *myo-2*p::mCherry (**c**) expression normalized to background at 48 h. **d** Representative images of animals fed OP50 and JUb19 at 72 h. **e, f** Quantification of *pals-5*p::GFP (**e**) and *myo-2*p::mCherry (**f**) expression normalized to background at 72 h. **b, c, e, f** 300 animals were quantified for each condition over three experimental replicates. In box-and-whisker plots, the central line marks the median, the box spans the interquartile range (25th–75th percentiles), and the whiskers extend to the smallest and largest values. Gray dots represent measurements for individual animals. Statistical significance was determined using an unpaired t-test, with significance levels denoted as: **** *p*<0.0001 (**b**, **e**), ns *p*=0.9692 (**c**), and *** *p*=0.0001 (**f**). **g** Representative images of *pals-5*p::GFP in specific tissues of OP50– and JUb19-fed animals, merged with DIC and red channel. Labeled structures: (1) intestine, (2) somatic gonad, (3) neurons, (4) epidermis, (5) rectal glands. **h** Representative images of IPR reporter induction in animals fed different *Stenotrophomonas* species. **i** Time course analysis of *pals-5*p::GFP induction in wild type and *zip-1(-)* animals. Dots represent percent of animals infected per experimental replicate. Statistical significance was determined using an unpaired t-test at each of the five timepoints, with significance levels denoted as: * *p*=0.0251, all remaining comparisons are not significant (ns), *p*>0.9999**. j.** Representative images of *pals-5*p::GFP induction of *pals-5*p::GFP in wild-type and *zip-1(-)* animals. **a, d, h, j** GFP images show *pals-5*p::GFP in cyan. **a, d, g, h, j** Composite images consist of merged green channel (*pals-5*p::GFP) shown in cyan, red channel (*myo-2*p::mCherry) shown in magenta and DIC images. Scale bars: 100 µm (**a, d, g, h, j**)

In contrast to canonical IPR-inducing intracellular pathogens such as Orsay virus and microsporidia, which induce *pals-5*p::GFP expression primarily in the intestine, JUb19-treated animals displayed reporter induction across multiple tissues (Fig. 1g)^14,19^. By 72 h, GFP expression was observed in the intestine, epidermis, neurons, somatic gonad, and rectal glands, suggesting a systemic transcriptional response rather than a localized one (Fig. 1g). Of note, weak expression was also observed in the posterior intestine and rectal glands on control OP50 plates. To determine whether this broad transcriptional activation is reflected at the protein level, we next examined the PALS-5::GFP translational reporter under the same conditions. In contrast to the transcriptional reporter, GFP signal was detected only in a subset of JUb19-treated animals and remained restricted to the cytosol of one or a few intestinal cells, with no detectable signal in other tissues (Supplementary Fig.1). This restricted pattern may reflect post-transcriptional regulation or decreased protein stability that limits PALS-5 accumulation outside the intestine despite transcriptional induction.

To test whether this response was unique to JUb19, we examined additional *Stenotrophomonas* isolates^22^. All tested *Stenotrophomonas* species and strains, including *Stenotrophomonas maltophilia* strains K279a, JV3, and JCMS; *Stenotrophomonas rhizophila* JUb23; and *Stenotrophomonas* sp. strains MYb57 and MYb240, also induced *pals-5*p::GFP expression (Fig. 1h), suggesting that IPR reporter induction is a conserved feature of the *Stenotrophomonas* genus^22^. Notably, qualitative differences in GFP intensity were observed, with pathogenic *S. maltophilia* strains appearing to induce stronger expression than non-pathogenic strains.

Previous studies showed that the bZIP transcription factor ZIP-1 functions as a key regulator of *pals-5*p::GFP expression in response to canonical IPR triggers^19,23,24^. We therefore asked whether JUb19 engages this same transcription factor. When animals were exposed to JUb19 for 48 h, we did not observe an obvious reduction in reporter expression in *zip-1* mutants compared to wild type (Supplementary Fig. 2). Because ZIP-1 regulates only the early phase of *pals-5* expression under certain conditions, we next performed a time-course assay to examine early reporter activation^19^. We monitored *pals-5*p::GFP expression in wild-type and *zip-1* mutant animals starting 3 h after transfer from OP50 to JUb19, when reporter expression first becomes detectable in wild type, and continuing to 5 h at 30 min intervals (Fig. 1i, j). Differences between genotypes were most apparent at 3 h post-transfer, when a smaller fraction of *zip-1*(-) animals expressed GFP compared to wild type (Fig. 1j, Supplementary Table 1). This difference decreased over time, and by 5 h, reporter expression converged such that the two genotypes were no longer detectably different (Fig. 1i, j, Supplementary Table 1). Although many individual timepoints did not reach statistical significance, the overall trend is consistent with ZIP-1 contributing to early JUb19-induced IPR activation, with later *pals-5*p::GFP expression sustained through ZIP-1–independent mechanisms^19^.

### JUb19-derived RNA induces *pals-5*p::GFP expression

Because the IPR is activated by obligate intracellular pathogens and does not overlap with immune programs triggered by extracellular bacterial pathogens, we hypothesized that JUb19 might invade intestinal cells and represent the first bacterial intracellular pathogen of *C. elegans*^11,13,14^. To test this, we used a *C. elegans* strain expressing *pept-1*p::PGP-1::GFP, which labels the apical membrane of intestinal cells and outlines the intestinal lumen^25,26^, and compared lumen boundaries with the distribution of FISH-stained bacteria (Fig. 2a). Animals were fed JUb19 or OP50 and collected at the L4, early adult, and gravid adult stage (Fig. 2a). While at the L4 stage we did not find noticeable intestinal colonization, minimal colonization was observed in early adults (Supplementary Fig. 3). In contrast, gravid adults displayed robust colonization throughout the intestinal lumen, particularly near the rectum, a pattern that was not observed in OP50-fed controls (Fig. 2a). These findings indicate that JUb19 remains confined to the intestinal lumen and does not act as an intracellular pathogen to induce the IPR.

**Figure 2:**
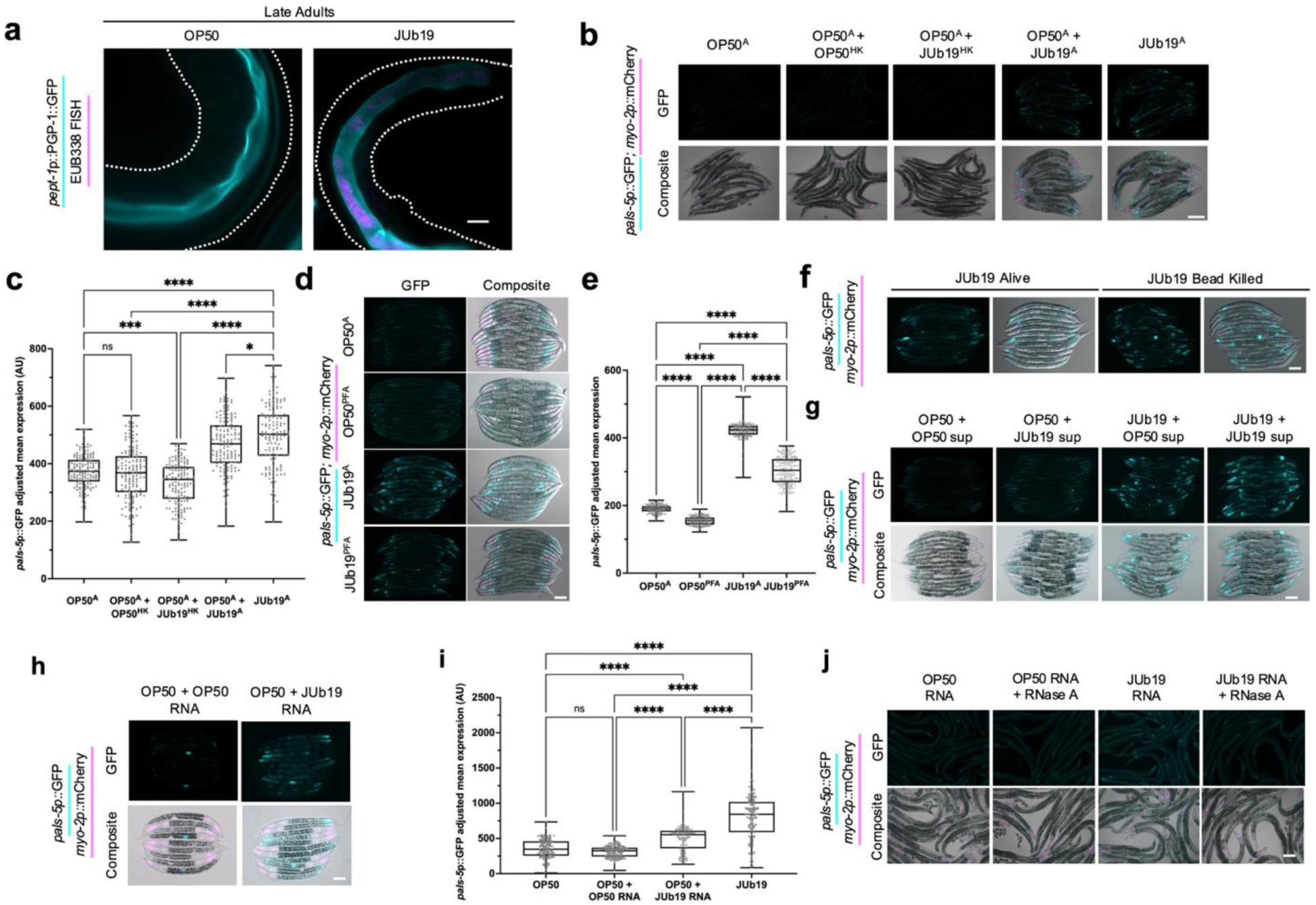
*S. indicatrix* JUb19 colonizes the *C. elegans* intestinal lumen and induces *pals-5*p::GFP via a heat-sensitive molecular compound. **a** Composite images of *pept-1*p::PGP-1::GFP intestinal apical membrane marker (cyan) and JUb19 labeled with EUB338 red fluorescent probe (magenta). Dotted white lines indicate the outline of each animal. Scale bar = 100 μm. **b** Representative images of IPR reporter induction following bacterial heat inactivation. **c** Quantification of *pals-5*p::GFP expression normalized to background fluorescence; 300 animals were analyzed per condition across three experimental replicates. Statistical significance was determined using Kruskal-Wallis test; OP50^A^ vs OP50^A^+OP50^HK^ *p*>0.9999, OP50^A^ vs. OP50^A^+JUb19^HK^ *p*=0.0003, OP50^A^ vs. JUb19^A^ *p*<0.0001, OP50^A^+OP50^HK^ vs. JUb19^A^ *p*<0.0001, OP50^A^+JUb19^HK^ vs. JUb19^A^ *p*<0.0001, JUb19^A^ vs. OP50^A^+JUb19^A^ *p*=0.0240. **(b, c)** A = alive, HK = heat-killed. **d** Representative images of animals fed chemically inactivated JUb19 and controls. **e** Quantification of *pals-5*p::GFP expression in animals fed chemically inactivated JUb19 and controls. Statistical significance was determined using a one-way ANOVA, Tukey’s test; p<0.0001 for all comparisons. **f** Representative images of IPR reporter induction following treatment with live and mechanically disrupted JUb19. **g** Representative images of IPR reporter induction following treatment with JUb19 supernatant. **h** Representative images of IPR reporter induction following treatment with OP50– and JUb19-derived RNA. **i** JUb19 RNA supplementation assay – IPR reporter expression quantification normalized to background fluorescence. Statistical significance was determined using Kruskal-Wallis test for multiple comparisons. *p*-values for **(i)** OP50 vs OP50+OP50 RNA *p*>0.9999, all remaining comparisons are *p*<0.0001. **j** Representative images of IPR reporter induction following treatment of OP50 and JUb19 RNA degraded by RNase A**. c, e, i** In box-and-whisker plots, the central line marks the median, the box spans the interquartile range (25th–75th percentiles), and the whiskers extend to the smallest and largest values. Gray dots represent measurements for individual animals. **b, d, f, g, h, j** Composite images consist of merged green channel (*pals-5*p::GFP) shown in cyan, red channel (*myo-2*p::mCherry) shown in magenta, and DIC images. Scale bars: 100 µm (**b, d, f, g, h, j**)

Next, we asked whether JUb19 viability is required for IPR reporter induction. We first inactivated the bacteria by heat treatment. Animals fed live JUb19 exhibited robust GFP induction, whereas those exposed to heat-killed JUb19 (mixed with live OP50) showed fluorescence comparable to OP50-fed controls (Fig. 2b, c, Supplementary Table 1), suggesting that either bacterial viability is required or the active factor is heat-labile. To distinguish between these possibilities, we next tested alternative inactivation methods. Paraformaldehyde (PFA)-treated JUb19 still induced *pals-5*p::GFP above OP50 controls, although induction was reduced relative to live JUb19 (Fig. 2d, e, Supplementary Table 1). Mechanical lysis of JUb19 also preserved its ability to induce IPR reporter expression (Fig. 2f). These data indicate that JUb19 does not need to be alive to trigger IPR reporter induction.

Previous work has shown that microbiome members can secrete factors that enhance *C. elegans* resistance to pathogens^7^. To test whether JUb19 activates the IPR via a secreted component, we supplemented plates seeded with either OP50 or JUb19 with sterile-filtered supernatants from JUb19 or OP50 cultures (Fig. 2g). After 48 h, animals grown on OP50 supplemented with JUb19 supernatant showed no detectable change in *pals-5*p::GFP expression relative to those exposed to OP50 supernatant alone (Fig. 2g). These findings suggest that JUb19 does not induce the IPR through soluble metabolites released into the surrounding medium.

Because our findings indicate that JUb19-induced IPR reporter activation depends on a heat-labile, non-secreted component, we hypothesized that bacterial RNA contributes to this activity (Fig. 2h). To test this, we supplemented bacterial lawns with total RNA purified from either JUb19 or OP50 and assessed *pals-5*p::GFP expression after 48 h (Fig. 2h). Because this initial exposure resulted in only minimal reporter induction, we repeated RNA supplementation at later developmental stages. Exposure to JUb19-derived RNA at both the L1 and L4 stages yielded a modest but detectable increase in GFP expression in adults, although induction remained lower than that observed with live JUb19 (Supplementary Fig. 4, Supplementary Table 1). A third supplementation at the early adult stage further increased *pals-5*p::GFP expression in JUb19 RNA-treated animals relative to both live OP50 and OP50-derived RNA controls (Fig. 2h-j, Supplementary Table 1).

Because double-stranded RNA (dsRNA) has previously been identified as an IPR trigger during Orsay virus infection through sensing by the RIG-I-like receptor DRH-1^20^, we next tested whether *pals-5p*::GFP induction by JUb19-derived RNA similarly requires DRH-1^20^. We found that reporter induction remained intact in *drh-1* mutants, suggesting that JUb19-derived RNA activates the IPR independently of canonical dsRNA sensing (Fig S5) To further evaluate the contribution of RNA, we treated JUb19 RNA preparations with RNase A, which preferentially degrades single-stranded RNA (ssRNA). RNase A treatment abolished reporter induction, suggesting that ssRNA contributes to JUb19-mediated IPR activation (Fig. 2j). Together, these findings indicate that JUb19 induces the IPR independently of intracellular colonization and bacterial viability and, at least in part, through an RNA-based mechanism.

### JUb19 triggers a distinct transcriptional response

We next evaluated whether JUb19 feeding influences endogenous mRNA expression of IPR genes using qRT-PCR, including *pals-5*, several IPR genes of unknown molecular function that are strongly induced by microsporidia infection (*F26F2.1, F26F2.3, F26F2.4*), and two genes encoding cullin-RING ubiquitin ligase components (*skr-5* and *cul-6*)^13,27,28^. Consistent with the increased *pals-5*p::GFP signal observed in JUb19-fed animals, we detected significant *pals-5* mRNA induction at 48 h compared with OP50-fed controls (Fig. 3a, Supplementary Table 1), with L4-stage animals exhibiting an average ∼5-fold increase. In contrast, no significant induction was observed for the other IPR genes at this time point (Fig. 3a, Supplementary Table 1). By 72 h, *pals-5* mRNA levels further increased to an average ∼10-fold induction relative to OP50-fed animals (Fig. 3b, Supplementary Table 1). *F26F2.1, F26F2.3, F26F2.4*, and *cul-6* expression levels remained similar to OP50 controls, whereas *skr-5* displayed a relatively robust ∼4-fold induction at 72 h (Fig. 3b, Supplementary Table 1). Overall, JUb19 elicits a relatively modest and partial IPR transcriptional signature, especially when compared with the broad IPR upregulation typically reported for microsporidia infection^13^.

**Figure 3:**
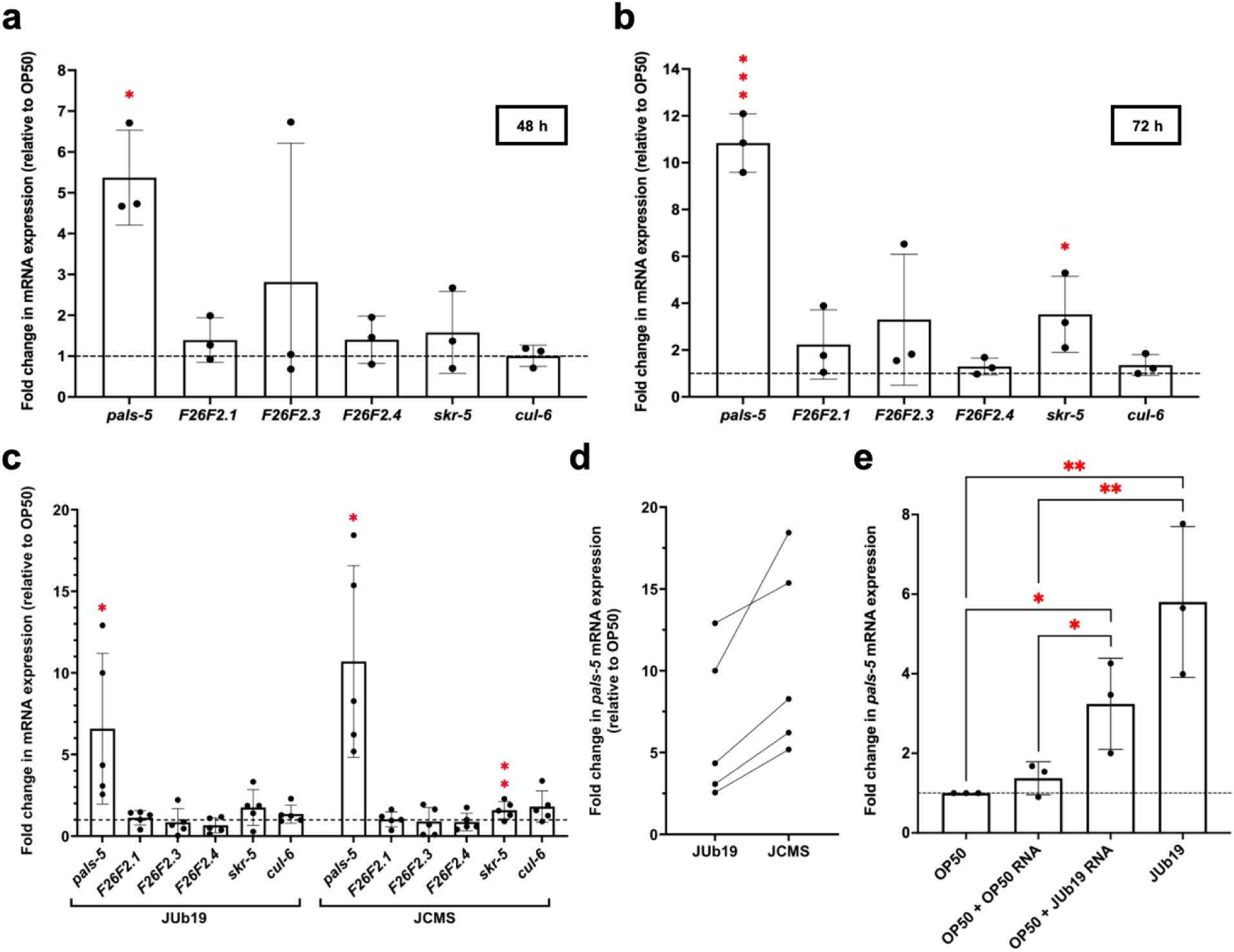
JUb19 induces expression of a subset of IPR genes. **a-c** qRT-PCR analysis of IPR gene expression in animals fed JUb19 (**a-c**) and JCMS (**c**) compared to OP50 control at 48 h (**a**), 72 h (**b**), or at L4 stage (**c**). **d** Stick and symbol graph showing the trend of *pals-5* mRNA expression between JUb19 and JCMS. **e** qRT-PCR analysis of *pals-5* expression following bacterial RNA supplementation. **a-d** Expression levels indicate fold changes relative to OP50 control. Values for each of the three experimental replicates are indicated with black dots. Bars indicate mean expression levels, with error bars representing standard deviation. Red asterisks above the bars indicate a statistically significant difference compared to the OP50 control; non-significant differences are not shown. Statistical significance was determined using a one-tailed t-test. *p*-values in (a): *pals-5 p*=0.0156, *F26F2.1 p*=0.1399, *F26F2.3 p* = 0.2032, *F26F2.4 p*=0.1464, *skr-5 p*=0.1863, *cul-6 p*=0.4833. *p*-values in (b): *pals-5 p*=8.41E05, *F26F2.1 p*=0.1102, *F26F2.3 p*=0.1139, *F26F2.4 p*=0.1112, *skr-5 p*=0.0273, *cul-6 p*=0.1181. *p*-values in (c): JUb19 – *pals-5 p*=0.01345, *F26F2.1 p*=0.2748, *F26F2.3 p*=0.3523, *F26F2.4 p*=0.0680, *skr-5 p*=0.0823, *cul-6 p*=0.0978; JCMS – *pals-5 p*=0.0030, *F26F2.1 p*=0.4390, *F26F2.3 p*=0.4478, *F26F2.4 p*=0.2715, *skr-5 p*=0.0193, *cul-6 p*=0.0588; *p*-values in (e): OP50 v. JUb19 RNA *p*=0.01389424, OP50 RNA v. JUb19 RNA *p*=0.02840618, OP50 + OP50 RNA v. OP50 + JUb19 RNA *p*=0.02840618, OP50 + OP50 RNA v. JUb19 *p*=0.00830299.

Because our GFP analyses suggested that more pathogenic *Stenotrophomonas* isolates may induce the IPR more strongly, we next compared qRT-PCR expression profiles of IPR genes in animals fed JUb19 versus the more pathogenic strain JCMS (Fig. 3c, Supplementary Table 1). Consistent with the stronger reporter expression, we found that *pals-5* levels showed a consistently higher trend in animals fed JCMS, although this difference did not reach statistical significance (Fig. 3c, d, Supplementary Table 1). The expression of other IPR genes was similar between strains and was not induced relative to the OP50 control, except for *skr-5* (Fig. 3c, Supplementary Table 1). Taken together, our analyses suggest that increasing *Stenotrophomonas* pathogenicity may be associated with increased *pals-5* expression (Fig. 3d, Supplementary Table 1).

Finally, to directly connect these transcriptional changes to the RNA-based activation observed in our reporter assays, we quantified *pals-5* expression levels following three sequential rounds of JUb19-derived RNA supplementation under the same conditions used in the IPR reporter expression analysis (Fig. 3e, Supplementary Table 1). As expected, animals fed live JUb19 displayed strong *pals-5* induction relative to live OP50. Supplementation with JUb19-derived RNA produced a modest but statistically significant increase in *pals-5* expression compared with both live OP50 and OP50-derived RNA, which showed only baseline or minimal induction (Fig. 3e, Supplementary Table 1). In summary, our qRT-PCR data are consistent with the reporter analysis and support a model in which *Stenotrophomonas* species induce IPR gene expression selectively, with response magnitude scaling with bacterial pathogenicity, and in which JUb19-derived RNA contributes, at least in part, to *pals-5* activation.

To characterize the global transcriptional response to JUb19, we performed RNA-seq on L4 animals fed JUb19 or OP50. JUb19-fed animals displayed a gene expression profile that diverged from those reported for canonical IPR triggers (Fig. 4a, b). Among 80 canonical IPR genes, only *pals-14* was strongly upregulated in JUb19-treated animals (Supplementary Table 2), consistent with its recently described role in defense against microsporidia^17^. Notably, *pals-5* was not detected as significantly upregulated, likely reflecting relatively modest induction and limited sequencing depth. Instead, the most highly induced genes (log₂ fold change > 2) included genes annotated or associated with innate immunity or pathogen responses (*lys-10, ilys-3, ilys-2, clec-70, asp-12*)^29–34^, fatty acid metabolism (*W03F9.4, ech-9, fat-7*)^30,35^, and cellular transport, transcriptional regulation, or xenobiotic/metabolic processes (*Y41C4A.32, arf-1.1, fkh-4, ceh-40, cyp-13B2*)^35–39^ (Supplementary Table 2).

**Figure 4:**
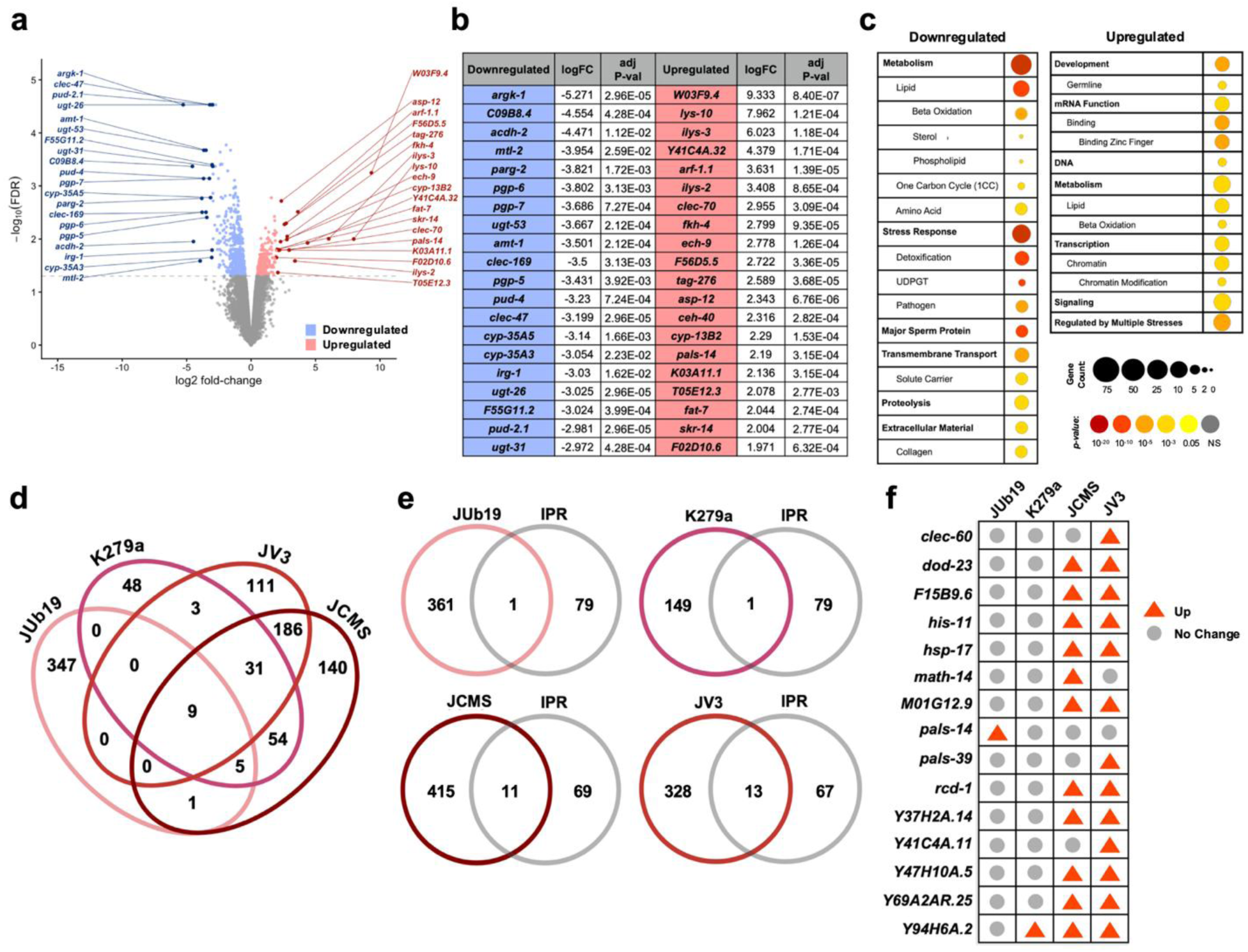
Analysis of transcriptional changes in worms exposed to JUb19. **a** Volcano plot of differently expressed genes in JUb19-fed worms vs. OP50 control. Each point on the plot represents a single gene. Significantly up– and downregulated genes (p < 0.05) are represented in light red and blue, respectively. Top 20 up– and downregulated genes with log2FC ≥ 2 are shown in dark red and blue, respectively. Genes with no change in expression are shown in grey. **b** Table showing top 20 up– and downregulated genes and the corresponding log2FC value. **c** Bubble plots of enriched categories for downregulated and upregulated genes following JUb19 treatment. Bubble area denotes category size (gene count), and color reports Bonferroni-adjusted *p*-values obtained from minimum hypergeometric tests. **d** Venn diagram of overlap between differentially expressed genes in worms exposed to JUb19 and other *Stenotrophomonas* strains. **e** Venn diagrams depict the comparison between upregulated genes in animals fed JUb19 and other *Stenotrophomonas* strains compared to the 80 canonical IPR genes. **f** Table of IPR genes upregulated following each analyzed *Stenotrophomonas* treatment.

To gain insight into the biological processes and cellular structures represented by these transcriptional changes, we used WormCat, a functional categorization tool optimized for *C. elegans* genomic data (Fig. 4c, Table 4). Upregulated genes were enriched for categories linked to responses to multiple stressors, mRNA binding and transcriptional regulation, and β-oxidation (Fig. 4c). In contrast, downregulated genes showed enrichment for stress-response pathways, particularly detoxification and pathogen defense, as well as metabolic processes related to lipid, amino acid, and one-carbon metabolism (Fig. 4c). Additional downregulated genes were grouped into categories associated with major sperm proteins and extracellular matrix/collagen (Fig. 4c). Thus, JUb19 exposure induces a mixed transcriptional program combining immune and metabolic signatures while suppressing specific stress– and reproduction-related pathways. To complement the WormCat analysis, we performed Gene Set Enrichment Analysis (GSEA) using curated immune and stress-related gene sets from published *C. elegans* transcriptional datasets (Supplementary Fig. 6, Supplementary Table 6). This analysis supported the conclusion that JUb19 does not broadly activate the IPR program, but instead shows selective enrichment of specific pathogen, stress, and metabolism-associated genes (Supplementary Fig. 6, Supplementary Table 6)

To place the JUb19 transcriptional response in the broader context of *Stenotrophomonas*-host interactions, we compared our RNA-seq dataset with previously published transcriptional profiles from animals exposed to three *S. maltophilia* strains: the non-pathogenic strain K279a and the pathogenic strains JV3 and JCMS^22^. At the global level, comparison of upregulated genes revealed limited overlap: only nine genes were induced across all four conditions, and JUb19 shared relatively few upregulated genes with each individual *S. maltophilia* strain (Fig. 4d). In contrast, overlap was higher between the two pathogenic strains, JV3 and JCMS, which shared 186 induced genes (Fig. 4d).

We next focused on canonical IPR genes by analyzing, for each bacterial condition, which of the 80 canonical IPR genes were upregulated (Fig. 4e, f). Unlike *pals-14,* which was upregulated in our dataset, all three *S. maltophilia* strains induced *Y94H6A.2*, an IPR gene of unknown function. Notably, this was the only IPR gene upregulated by the non-pathogenic K279a, whereas the pathogenic strains JV3 and JCMS induced an additional 12 and 10 IPR genes, respectively (Fig. 4e, f). Nine of these, together with Y94H6A.2, were upregulated in both JV3– and JCMS-treated animals (Fig. 4e, f). Thus, although the overall overlap between JUb19 and *S. maltophilia* transcriptional responses is modest, analysis of canonical IPR genes reveals a consistent pattern: more pathogenic *Stenotrophomonas* strains are associated with broader and stronger IPR gene activation than less pathogenic or non-pathogenic strains, with JUb19 eliciting a comparatively limited IPR induction.

### JUb19 provides resistance to obligate intracellular pathogens

Although robust IPR activation has been linked to enhanced resistance to obligate intracellular pathogens^16,18,19,40–42^, our transcriptomic analysis indicates that JUb19 only modestly engages the canonical IPR and instead induces a distinct transcriptional program, raising the question of whether this response confers protection against infection. We first assessed susceptibility to viral infection. At the L4 stage, JUb19-fed animals showed a lower frequency (∼38%) of Orsay virus infection than OP50-fed controls (∼63%) (Fig. 5a, Supplementary Table 1). We next tested whether JUb19 similarly protects against microsporidian *N. parisii*. JUb19-fed animals exhibited significantly reduced infection compared to OP50-fed controls. When L4-stage animals were infected in liquid and scored as adults, approximately 5% of JUb19-fed animals were infected versus approximately 20% of OP50-fed animals (Fig. 5b, Supplementary Table 1). In a complementary experiment using the standard 30-hour *N. parisii* infection protocol initiated at the L1 stage, JUb19-fed animals displayed significantly lower pathogen loads than OP50-fed controls (Fig. 5c, Supplementary Table 1). In summary, these results demonstrate that JUb19 exposure confers protection against evolutionarily and molecularly distinct obligate intracellular pathogens. To determine whether reduced pathogen burden could be explained by differences in food intake or waste elimination, we compared pharyngeal pumping and defecation rates between JUb19-and OP50-fed animals. Neither pharyngeal pumping nor defecation rates differed significantly between conditions (Supplementary Fig. 7, Supplementary Table 1). These findings indicate that JUb19-mediated protection is not due to reduced feeding or altered gut transit and is unlikely to result from decreased pathogen exposure.

**Figure 5:**
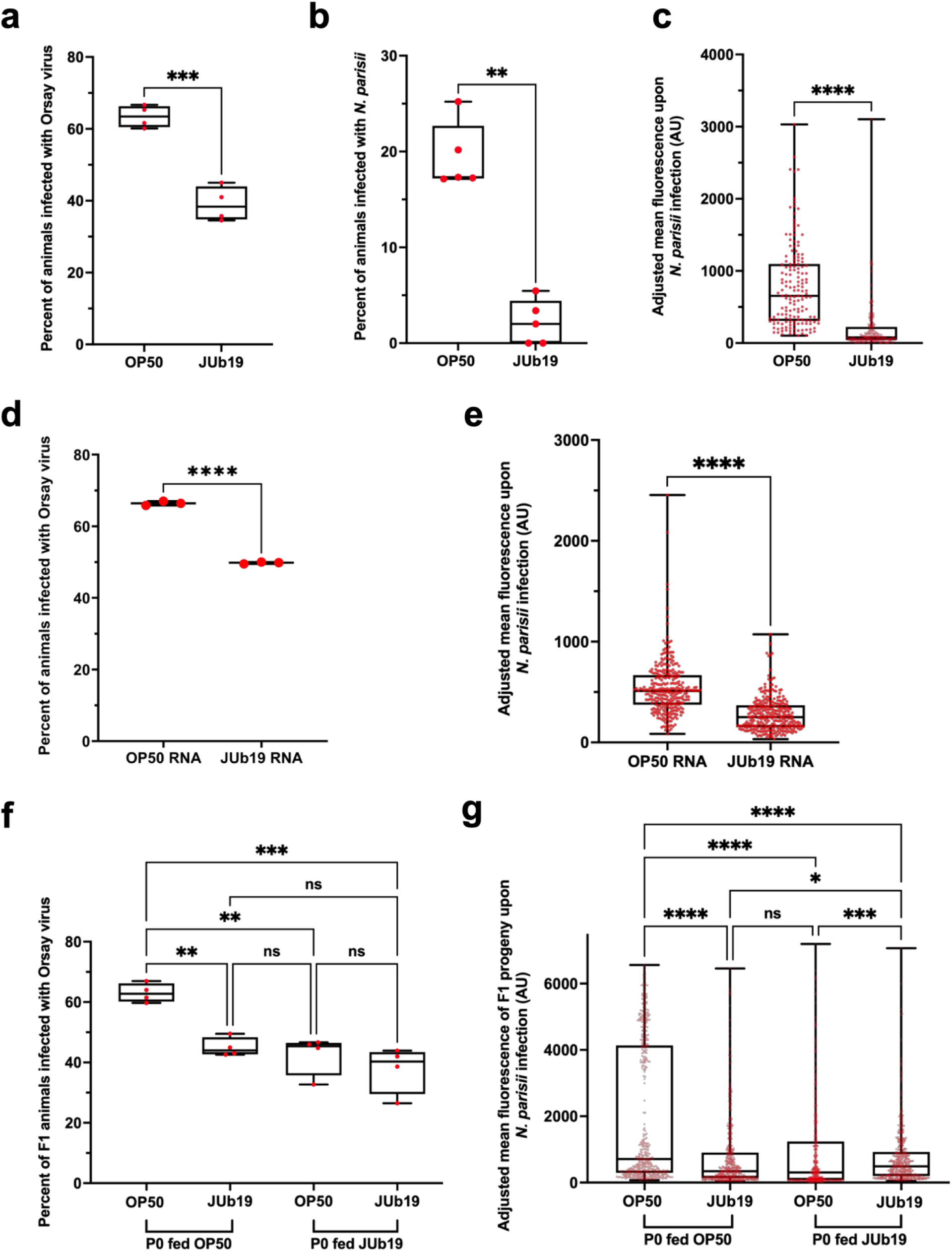
JUb19 protects *C. elegans* against obligate intracellular pathogens. **a** Fraction of animals infected with Orsay virus. **b** Fraction of animals infected with *N. parisii*. **c** Quantification of *N. parisii* mean FISH fluorescence signal normalized to body area. **a-c** 300 animals were analyzed per experimental replicate. Statistical significance was determined using an unpaired t-test; *p*-values: *p*=0.0004 **(a)**, *p*=0.0079 **(b),** *p*<0.0001 **(c). d** Fraction of animals infected with Orsay virus following RNA supplementation. 300 animals were analyzed in each experimental replicate. **e** Quantification of N. parisii mean FISH fluorescence signal normalized to body area. 100 animals were analyzed per experimental replicate. Statistical significance was determined using an unpaired t-test; *p*-values: *p*<0.0001 (**e, d**), *p*=0.0001 **f** Fraction of animals infected with Orsay virus in F1 generation. **g** Quantification of *N. parisii* mean FISH fluorescence signal normalized to body area in F1 generation. **f, g** 300 (**f**) and 150 (**g**) animals per condition per three experimental replicates were analyzed. Samples exposed to OP50 and JUb19 in the P0 generation are shown shaded in light blue and light yellow, respectively. Statistical significance was determined using the Kruskal-Wallis test for multiple comparisons. *p*-values for **(f):** OP50 (P0 – OP50) vs. OP50 (P0 – JUb19) *p*=0.0185, OP50 (P0 – OP50) vs. JUb19 (P0 – JUb19) *p*=0.0073, all other comparisons are non-significant (ns) *p*<0.9999. *p*-values for (**g**): JUb19 (OP50-P0) vs. JUb19 (JUb19-P0) *p*=0.0213, OP50 (P0-JUb19) vs. JUb19 (P0-JUb19) *p*=0.0003, remaining comparisons are either ns (*p*>0.999) or **** *p*<0.0001. **a-g** In box-and-whisker plots, the central line marks the median, the box spans the interquartile range (25th–75th percentiles), and the whiskers extend to the smallest and largest values. Red dots in **a, b, d, f** represent percent of animals infected per experimental replicate; red dots in **c, e, g** represent measurements for individual animals.

Because JUb19-derived RNA was sufficient to induce *pals-5* reporter and mRNA expression (Fig. 2h, Fig. 3e), we next asked whether RNA supplementation was also sufficient to confer protection against intracellular pathogens. Using the same supplementation strategy in which RNA was administered three times during development, we first assessed susceptibility to Orsay virus. JUb19 RNA supplementation consistently reduced viral infection frequency relative to animals supplemented with OP50 RNA (Fig. 5d, Supplementary Table 1). Furthermore, JUb19-derived RNA reduced *N. parisii* pathogen load (Fig. 5e, Supplementary Table 1). Together, these results suggest that JUb19-derived RNA is sufficient to at least partially recapitulate both IPR induction and pathogen protection associated with live JUb19 exposure.

Given previous work showing that *N. parisii* infection in the parental generation can confer inherited immunity to progeny^43^, and our observation of *pals-5*p::GFP expression in the somatic gonad, we next asked whether JUb19 exposure similarly promotes inherited immunity. To test this, we exposed parental (P0) animals to JUb19 and then assessed the susceptibility of F1 progeny to either Orsay virus or *N. parisii* infection after the progeny were raised on OP50 (Fig. 5f,g, Supplementary Table 1). Offspring of JUb19-exposed mothers showed significantly reduced Orsay virus infection frequency compared to progeny from parents maintained on OP50 alone, with protection levels comparable to those observed under continuous JUb19 exposure across both generations (Fig. 5f, Supplementary Table 1). A similar inherited protective effect was observed for *N. parisii* infection (Fig. 5g, Supplementary Table 1). Thus, JUb19 exposure in the parental generation is sufficient to confer inherited immunity against multiple intracellular pathogens, although its underlying mechanisms remain to be defined.

### JUb19 exposure is associated with impaired fitness in *C. elegans*

Constitutive immune activation in *C. elegans* is often accompanied by developmental defects, a phenotype we also observed in animals fed JUb19^21^. At the L4 stage, JUb19-fed animals showed a mild but statistically significant reduction in body length compared with OP50-fed controls (Fig. 6a, Supplementary Table 1). This size difference became more pronounced in adults, where JUb19-fed animals appeared visibly smaller and exhibited a significant reduction in body length (Fig. 6b, Supplementary Table 1). These observations indicate that prolonged exposure to JUb19 causes a measurable developmental delay, consistent with phenotypes associated with chronic IPR activation^16,21,40,42,44^.

**Figure 6:**
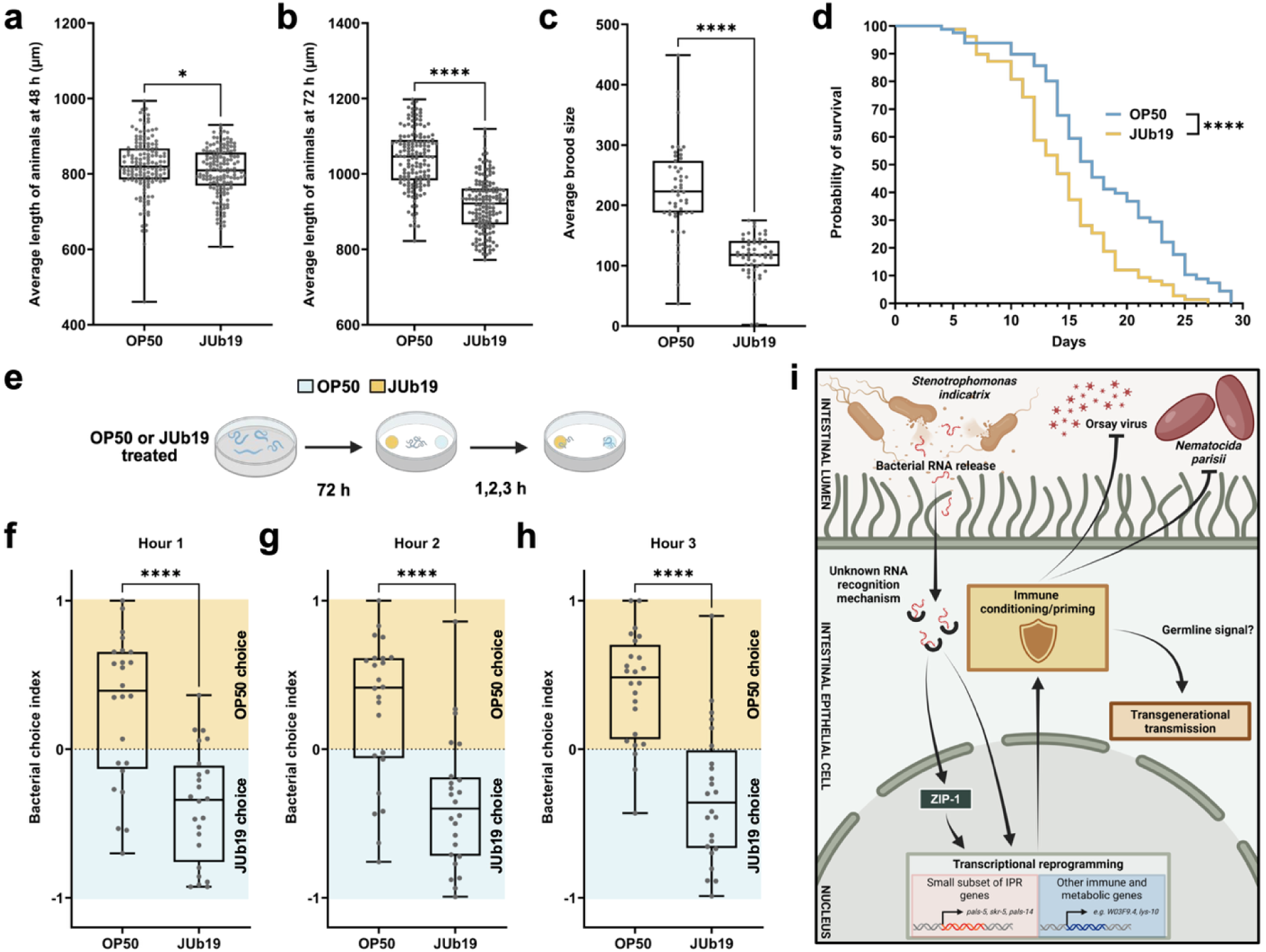
JUb19 impairs *C. elegans* fitness and influences dietary preference. **a, b** Analysis of body length of animals grown on OP50 and JUb19 at 48 h (**a**) and 72 h (**b**). 300 animals were analyzed per condition, per each of three experimental replicates. Box and whisker plots of measured length of OP50– and JUb19-fed animals at 48 h. **c** Brood size analysis of ten animals per condition per experimental replicate. **a-c** In box-and-whisker plots, the central line marks the median, the box spans the interquartile range (25th–75th percentiles), and the whiskers extend to the smallest and largest values. Gray dots represent measurements for individual animals. Statistical significance was determined using an unpaired t-test, *p*-values: * *p*=0.0406 (**a**), **** *p*<0.0001 (**b, c**). **d** Lifespan analyses of animals grown on OP50 and JUb19, 40 animals were analyzed per replicate, three replicates total. Statistical significance was determined using Log-rank (Mantel-Cox) test, *p*-value **** *p*<0.0001. **e** Schematic representation of food choice assay. Created with BioRender. **f-h** Analysis of bacterial choice index across three consecutive time points. Bacterial choice index for JUb19 was calculated as (# of animals on JUb19) – (# of animals on OP50)/(total). A positive index indicates a preference for JUb19 over OP50, while a negative index indicates a preference for OP50 over JUb19. In box-and-whisker plots, the central line marks the median, the box spans the interquartile range (25th–75th percentiles), and the whiskers extend to the smallest and largest values. Each dot represents a result from a single plate; eight plates were analyzed per each of three experimental replicates. Statistical significance was determined using an unpaired t-test, *p*-value across all three time points for OP50 vs. JUb19 *p*<0.0001. **i** Model of JUb19 and *C. elegans* interaction. Created with BioRender.com.

To determine whether this developmental delay reflects a broader fitness cost, we examined additional fitness parameters. JUb19-fed animals produced significantly fewer progeny than OP50-fed controls, indicating reduced reproductive output (Fig. 6c, Supplementary Table 1). Furthermore, lifespan analysis revealed a significant reduction in longevity in JUb19-fed animals (Fig. 6d, Supplementary Table 1). Together, these results suggest that JUb19-induced immune activation is coupled to a general decline in host fitness, encompassing impaired growth, smaller brood size, and shorter lifespan. In addition, these data may indicate that JUb19 is mildly pathogenic to *C. elegans*.

We next asked whether this fitness cost influences bacterial choice. In a two-choice assay, adult animals raised on either OP50 or JUb19 were transferred to plates containing adjacent lawns of both bacteria, and their distribution was scored at 1, 2, and 3 h time points after transfer (Fig. 6e, Supplementary Table 1). Animals previously exposed to JUb19 consistently avoided JUb19 and showed a strong preference for OP50 at all time points (Fig. 6f, Supplementary Table 1). By contrast, OP50-raised animals initially preferred JUb19, although this preference decreased slightly over time. These findings indicate that prior exposure to JUb19 promotes learned avoidance, potentially representing a behavioral adaptation to limit the fitness costs associated with JUb19-induced immune activation.

## Discussion

Our results revise the current view of the IPR as a program engaged only by intracellular infection^13,14,19,27^. We show that exposure to a natural microbiome bacterium is sufficient to partially induce the IPR and establish resistance to intracellular pathogens without intracellular invasion and before stable colonization. These findings indicate that microbiota can shape immune readiness prior to pathogen encounter.

Transcriptomic analyses further show that JUb19 does not induce the full IPR but instead triggers broader immune–metabolic transcriptional changes. This pattern suggests that pathogen resistance in JUb19-treated animals reflects a distinct, integrated response rather than a single defense pathway. Interestingly, several of the most strongly differentially expressed genes in JUb19-treated animals, including the upregulated genes *fat-7*, *lys-10*, *ilys-2*, and *ilys-3*, as well as the downregulated gene *argk-1*, were also reported to be differentially expressed in *C. elegans* fed *Microbacterium* sp. JUb74, a fat-rich diet that causes developmental and reproductive defects without IPR activation, although resistance to intracellular pathogens has not been examined in this context.^30^. Future studies will be needed to determine whether immune protection can be uncoupled from fitness costs, particularly if these phenotypes are governed by distinct genetic programs, or whether metabolic changes shared between JUb19– and *Microbacterium*-fed animals contribute to defense.

The observation that all tested *Stenotrophomonas* species induce IPR reporter expression indicates that this response is a genus-level feature of host interaction rather than a strain-specific effect. However, more pathogenic strains induce stronger and broader IPR activation, suggesting that IPR amplitude scales with bacterial pathogenicity^22^. In addition, the pathogenic *S. maltophilia* strain JCMS has been shown to evade the DAF-2/DAF-16 insulin signaling pathway^45^, a conserved antibacterial immune program in *C. elegans*^22,46,47^. These findings raise the possibility that bacterial induction of the IPR represents an evolutionary strategy in which activation of defenses against intracellular pathogens redirects host immune resources away from antibacterial defenses, imposes fitness costs, and thereby potentially favors bacterial persistence and pathogenicity.

Our data also identify bacterial RNA as a microbial cue that contributes to partial IPR induction and pathogen protection. While bacterial dsRNA has previously been proposed to induce interferon beta production in mammals and promote intestinal immune homeostasis^48^, our study provides functional evidence that bacterial RNA can directly promote resistance to intracellular infection. Furthermore, previous observations of bacterial RNA activity primarily involved responses directed toward the source organism, including bacterial avoidance in *C. elegans* and immune priming in plants and molluscs.^49–52^. In contrast, our findings show that bacterial RNA can modify host susceptibility and confer protection against unrelated intracellular infections caused by viral and microsporidian pathogens. including viral and microsporidian infection. However, the contribution of RNA to reporter induction and pathogen resistance appears to be partial compared to live JUb19, possibly because RNA acts together with additional bacterial factors or because its activity is reduced outside the native bacterial context.

JUb19 exposure additionally generates inherited protection against viral and microsporidian infection, demonstrating that microbiome-derived signals alone can establish transgenerational defensive states. Given the prevalence of microsporidia in natural *C. elegans* populations, such priming may provide adaptive benefits despite measurable fitness costs under laboratory conditions^11,15^. Taken together, our findings support a model in which dietary microbes tune intracellular pathogen resistance by coupling partial activation of a known immune response with systemic physiological remodeling, linking microbiome composition, host state, and infection outcome.

## Materials and Methods

### *C. elegans* maintenance

*C elegans* were maintained on Nematode Growth Medium (NGM) plates at 20 °C^53,54^; plates were seeded with *E. coli* OP50, *S. indicatrix* strain JUb19, or other *Stenotrophomonas* spp. unless stated otherwise. *C. elegans* list of strains is outlined in Supplementary Table 7.

### Synchronization of *C. elegans* via bleaching

Synchronization was achieved by transferring 25–30 L4 *C. elegans* to seeded NGM 2-4 plates for 96 h at 20 °C. Animals were collected with M9, pelleted (2,400 × g, 40 s), and resuspended in 3 mL of M9, and then 1 mL bleaching solution (1:1 mixture of 5% sodium hypochlorite and 5 M NaOH) was added. Eggs were washed five times with M9 and incubated in 3 mL of M9 at 20 °C for 18–24 h to yield L1-synchronized populations^55^.

### Bacterial maintenance

Single bacterial colonies grown on Luria–Bertani (LB) agar were inoculated into LB broth to generate overnight cultures. *E. coli* OP50 was grown at 37 °C, whereas JUb19 and other *Stenotrophomonas* strains were grown at 25 °C.

### Fluorescent reporter measurements for initial screen

For L4 and adult GFP expression analysis on JUb19, synchronized L1 larvae were exposed to OP50 or JUb19 at 20 °C for 48 or 72 h, fixed in 4% paraformaldehyde (PFA) for 7 min, and imaged on a Zeiss LSM 700 confocal microscope using software ZEN 2010. Mean fluorescence of *pals-5*p::GFP and *myo-2*p::mCherry was quantified per animal in FIJI (ImageJ)^56^, background-normalized, and pooled across three biological replicates (100 animals each; total n = 300).

### Assessment of ZIP-1 requirement for JUb19-induced IPR reporter activation

A time-course experiment was performed using a control strain and a *zip-1(jy13)* mutant, both carrying the *jyIs8* transgene. Synchronized populations (300 animals per strain) were cultured on OP50 for 48 h at 20 °C and then transferred to JUb19. GFP expression was assessed in 50 animals per condition beginning 3 h after transfer (the earliest time point at which *pals-5*p::GFP was detectable in the control), and subsequently every 30 min for an additional 2 h until the final time point at 5 h. Imaging and quantification were conducted on a Zeiss Axio Imager.M2 compound microscope equipped with a Zeiss Axiocam 820 mono camera and ZEN 3.8 software.

### Bacterial fluorescence in situ hybridization (FISH) analysis

Developmental rate of GK288 (*dkIs166[pept-1p*::*pgp-1::gfp]*) was assessed by staging L4 (63 h), young adult (72 h), and later adult (96 h) animals at 20°C, as this strain develops more slowly than N2 under the same conditions. Synchronized L1s were plated on 10 cm NGM seeded with OP50 or JUb19 and harvested at each time point. Animals were washed with M9 + 0.1% Tween, centrifuged (200 rcf, 30 s), fixed in 4% PFA, and processed for FISH: overnight hybridization at 46 °C in 100 µL buffer (900 mM NaCl, 20 mM Tris pH 7.5, 0.01% SDS) with Red-conjugated EUB338 (5’-GCTGCCTCCCGTAGGAGT-3’) universal FISH probe targeting 16s rRNA^57^, followed by a 1 h wash at 46 °C in wash buffer (900 mM NaCl, 20 mM Tris, 5 mM EDTA, 0.01% SDS) and a final PBS-T rinse. Imaging used a Zeiss Axio Imager.M2 compound microscope equipped with Zeiss Axiocam 820 camera and ZEN 3.8 software.

### Bacterial inactivation by heat

JUb19 and OP50 were cultured overnight at 37 °C and heat-inactivated at 60 °C for 60 min. Loss of viability was verified by spotting 10 µL of culture onto LB agar and incubating at 37 °C for 48 h; absence of colonies was taken as confirmation. Heat-killed preparations were then used to seed NGM plates and, to prevent developmental arrest, were supplemented with live *E. coli* OP50. Mixtures of heat-killed JUb19 and live OP50, along with live JUb19 and OP50 controls, were spotted onto NGM plates for GFP expression assays. Worms were maintained on these plates for 72 h at 20 °C, fixed in 4% PFA for 5 min, and imaged for representative fluorescence on a Zeiss LSM 700 confocal microscope operated via ZEN 2.1. For quantitative fluorescence analysis, animals were imaged using an ImageXpress Nano automated imaging system, and images were processed in FIJI (ImageJ).

### Chemical inactivation of bacteria

Bacterial strains were cultured overnight, harvested, and fixed with 0.5% PFA (prepared by mixing 15.5 parts culture with 0.5 parts 16% PFA to yield 0.5% final). Mixtures were incubated at 37 °C for 30 min, pelleted at 6,000 G for 5 min, and washed three times with LB to remove residual PFA. Loss of viability was verified by spotting 10 µL of culture onto LB agar and incubating at 37 °C for 48 h; absence of colonies was taken as confirmation. PFA-treated bacteria were then used to supplement NGM plates. Approximately 300 synchronized L1s were placed on these plates and maintained at 20 °C for 48 h. Animals were collected in M9 and anesthetized with 100 mM levamisole for fluorescence imaging and GFP quantification. Imaging was performed on a Zeiss Axio Imager.M2 compound microscope equipped with a Zeiss Axiocam 820 mono camera and ZEN 3.8 software. Image analysis was carried out in FIJI (ImageJ).

### Mechanical inactivation of bacteria

JUb19 was cultured overnight and mechanically lysed using 0.1-mm zirconia/silica beads (BioSpec Products). Disruption was performed on a Digital Disruptor Genie (Scientific Industries) at 3,000 rpm for 60 min at 4 °C. The resulting supernatant was collected and used to seed NGM plates. The absence of viable bacteria in the supernatant was verified as described above by spotting samples onto LB agar plates and confirming the absence of colony growth after incubation. Approximately 300 synchronized L1 larvae were transferred to each treatment plate and to plates seeded with live JUb19 as a control, and maintained at 20°C for 48 h. Animals were then collected in M9 and anesthetized with 100 mM levamisole for fluorescence imaging.

### Bacterial supernatant test

Overnight cultures of OP50 and JUb19 were grown in LB at 37 °C and 25 °C, respectively. 50 mL cultures were centrifuged at 4,300 rpm for 10 min, and the supernatants were collected and sterile-filtered using a 0.2 µm vacuum filtration system. Filtered supernatants were applied to NGM plates pre-seeded with OP50 or JUb19. Synchronized L1-stage *C. elegans* were transferred to these plates and allowed to develop to L4 – 48 h for OP50 and 50 h for JUb19 to account for the minor developmental delay with JUb19 feeding. Representative fluorescent images were acquired on a Zeiss Axio Imager.M2 compound microscope equipped with a Zeiss Axiocam 820 mono camera and ZEN 3.8 software. Image analysis was performed in FIJI (ImageJ).

### Bacterial RNA supplementation

Overnight LB cultures (50 mL each) of OP50 (37 °C) and JUb19 (25 °C) were harvested into TRI reagent (Molecular Research Center, Inc.), flash-frozen, and subjected to three freeze–thaw cycles. RNA was isolated by BCP phase separation (Molecular Research Center, Inc.), purified by isopropanol precipitation and ethanol washes, adjusted to ∼1000 ng/µL, and stored at –80 °C until further use. For RNA degradation, RNA samples were treated with RNase A by adding 2 µL RNase A to 50 µL RNA and incubating at 47 °C for 15 min. RNase-treated samples were then stored at –80 °C until use. Synchronized L1-stage *C. elegans* were plated onto OP50 lawns pre-primed with 100 µL of bacterial RNA (∼800 ng/µL); additional RNA was applied at 48 h. A final supplementation was performed at the gravid adult stage (72 h) in liquid: worms were collected from plates using M9, resuspended in purified RNA, and incubated for 1 h before imaging. Representative fluorescent images were acquired on a Zeiss Axio Imager.M2 compound microscope equipped with a Zeiss Axiocam 820 mono camera and ZEN 3.8 software, and images were analyzed in FIJI (ImageJ).

### *C. elegans* RNA isolation

Animals were rinsed from OP50– and JUb19-seeded plates with M9 buffer and transferred into TRI Reagent. Suspensions were incubated at room temperature for 30 min, thoroughly homogenized, and stored at −80 °C. Total RNA was isolated by BCP phase separation, then purified by isopropanol precipitation followed by a 75% ethanol wash. For RNA-seq, JUb19 fed conditions were collected 2h later than OP50 conditions to better match developmental differences across conditions. Samples received an additional cleanup with the RNeasy Universal Mini Kit per manufacturer’s instructions (Qiagen).

### qRT-PCR analysis

cDNA was synthesized from total RNA using the iScript cDNA Synthesis Kit (Bio-Rad) according to the manufacturer’s instructions. qRT-PCR was performed with iQ SYBR Green Supermix (Bio-Rad) on a CFX real-time PCR detection system (Bio-Rad). For each target, two technical replicates were run within each of three biological replicates. Gene expression was normalized to *snb-1*, which does not change expression during IPR activation. Primer sequences are listed in Supplementary Table 8. Data analysis was carried out in GraphPad Prism using the Pfaffl method^58^, and one-tailed t-tests were used to assess differences between samples.

### RNA sequencing analysis

cDNA libraries were prepared and sequenced (paired end) by Genewiz Azenta. Sequencing reads were aligned to the *C. elegans* reference genome (WormBase release WS235) using Rsubread (run in RStudio). Gene-level counts were normalized and analyzed for differential expression with the limma-voom pipeline on the Galaxy platform (usegalaxy.org). Genes with <1 count per million (CPM) across all samples were excluded, and differentially expressed genes were defined at an adjusted p-value < 0.05. Differential expression results were visualized as a volcano plot showing log₂ fold change versus −log₁₀ (adjusted p-value). Functional enrichment was assessed using Gene Set Enrichment Analysis (GSEA v4.3.2)^59^ against curated immune gene sets, with normalized enrichment scores (NES), nominal p-values, and false discovery rate (FDR) q-values reported (Supplementary Table 6). Category-level enrichment of differentially expressed genes (DEGs) was performed with WormCat 2.0 (wormcat.com)^60^ and visualized as category-level enrichment plots. Overlap with published immune-response datasets was summarized with Venn diagrams generated using Intervene (https://intervene.readthedocs.io/en/latest/index.html), highlighting shared and unique DEGs (see Fig. 4).

### Orsay infection

To increase infection sensitivity, Orsay virus experiments were performed using WM27 *[rde-1(ne219)]* animals, which are defective in antiviral RNA interference. Synchronized L1 animals were grown to the L4 stage on plates seeded with OP50 or JUb19 for 48 h or 50 h, respectively, to account for the slight developmental delay associated with JUb19 feeding. L4 animals were then exposed to Orsay virus mixed with their respective food source (OP50 or JUb19), maintaining the same bacterial condition as during initial plating, and incubated for 24 h at 20 °C. After incubation, animals were fixed in 4% PFA for 15 min and stained overnight at 46 °C with Orsay1 Cal Fluor Red 610 (5′-GACATATGATGCCGAGAC-3′) and Orsay2 Cal Fluor Red 610 (5′-GTAGTGTCATTGTAGGCACC-3′) probes, targeting Orsay virus RNA1 and RNA2, respectively^57,61^. Infection percentages were scored on a Zeiss Axio Imager.M2 compound microscope equipped with a Zeiss Axiocam 820 mono camera and ZEN 3.8 software. For each of three experimental replicates, 300 animals per condition were assessed for infection. For RNA supplementation experiments, RNA was supplemented at the L1 (1h), L4 (48 h), and adult (72 h) stages. Infection and FISH staining were performed as described above.

### Microsporidia infection

To quantify the proportion of infected animals, synchronized L1 *C. elegans* (N2) were grown on OP50– or JUb19-seeded NGM plates (1,200 animals per plate) at 20 °C for 48 h, then washed off plates with M9 buffer into 1.5 mL microcentrifuge tubes, pelleted, and resuspended to 100 µL. *N. parisii* spores (3 million per sample) were added in M9 to a total volume of 500 µL, and tubes were rotated at 25 °C for 15 min. Worms were then transferred onto fresh NGM plates (250 µL per plate; two plates per condition), allowed to dry, and incubated at 25 °C for 24 h. Animals were fixed and stained with MicroB-CF610 (5′-CTCTCGGCACTCCTTCCTG-3′), a Cal Fluor Red 610-conjugated probe targeting ribosomal RNA of microsporidia^57^, and at least 100 animals per replicate were scored for infection on a Zeiss Axio Imager.M1 compound microscope using AxioVision 4.8.2. Five experimental replicates were performed.

For pathogen load measurements, synchronized L1 wild-type *C. elegans* were mixed with food (OP50 or JUb19) and *Nematocida parisii* spores (300,000 per sample), and this mixture was added to unseeded NGM plates and incubated at 25 °C for 30 h. Animals were fixed in 4% PFA for 30 min and stained overnight at 46 °C with MicroB-CF610^57^. Imaging was performed on a Zeiss Axio Imager.M2 compound microscope equipped with a Zeiss Axiocam 820 mono camera and ZEN 3.8 software, and fluorescence was quantified in FIJI (ImageJ). For each condition in each of three biological replicates, 100 animals were analyzed; mean fluorescence per animal was background normalized. For assessing infection following RNA supplementation. RNA was supplemented at 1 h, 8 h, and 29 h within the 30 h infection period.

### Pharyngeal pumping analysis

Pharyngeal pumping, as a measure of feeding rate, was quantified in wild-type animals and in *eat-2(ad465)* mutants (strain DA465)^62,63^, which exhibit a reduced pumping rate. L4 animals grown at 20 °C on OP50 or JUb19 were transferred to fresh lawns of the same bacterial condition. Pharyngeal pumping was quantified by counting terminal bulb contractions over three consecutive 20 s intervals; the mean number per 60 s was recorded as the feeding rate. For each condition, 40 animals were scored in each of three experimental replicates.

### Defecation rate analysis

Defecation rate assays were performed in wild-type animals and in mutants with altered defecation frequency, including JT89 (*dec-2(sa89)*), which exhibits a slower defecation rate, and JT296 (*dec-7(sa296)*)^64^, which exhibits a faster defecation rate. Animals were maintained on OP50 or JUb19 at 20 °C. Gravid adults were assessed for defecation cycle length, defined as the interval (seconds) between successive defecation events. For each strain and condition, five animals were monitored for five complete cycles in each of three experimental replicates, and the mean cycle time per individual was recorded^64^.

### Inherited immunity assay

Parental (P0) populations were raised for 96 h at 20 °C on NGM plates seeded with either OP50 or JUb19. Synchronized F1 progeny were obtained by bleaching. F1 progeny derived from each P0 condition were then plated onto OP50– or JUb19-seeded plates (yielding four conditions: P0-OP50 to F1-OP50, P0-OP50 to F1-JUb19, P0-JUb19 to F1-OP50, P0-JUb19 to F1-JUb19) and maintained at 20 °C to the L4 stage (48 h for OP50 treatments and 50 h for JUb19 treatments).

For Orsay infection at L4, F1 animals were infected with Orsay virus and processed for FISH quantification exactly as described in the Orsay virus infections section. For microsporidia assays, synchronized F1 L1 animals from each P0 condition were infected with *N. parisii* for 30 h following the microsporidia pathogen load assay protocol described above (same spore dose, temperature, staining, and scoring).

### Body length measurements

Animals carrying the IPR reporter (strain ERT054, *jyIs8[pals-5p::gfp; myo-2p::mCherry]*) were synchronized, and L1 larvae were placed on control OP50 or JUb19 and maintained at 20 °C for 48 h or 72 h. Animals were collected at each time point and fixed in 4% PFA. Body length was measured in FIJI (ImageJ). In total, 300 animals were analyzed (100 animals per condition in each of three experimental replicates).

### Longevity and brood size analysis

Synchronized N2 populations were grown on OP50 or JUb19 for 48 h at 20 °C. At 48 h, 40 animals (20 per condition) were transferred to 3.5 cm NGM plates seeded with OP50 or JUb19 and moved daily until death. Longevity and egg laying were monitored daily. Four biological replicates of longevity analysis were analyzed using the Survival function in GraphPad Prism 10, and statistical significance was assessed with the log-rank (Mantel–Cox) test.

### Bacterial preference test analysis

For plate preparation, single colonies of JUb19 were inoculated in LB medium and grown overnight at 25 °C, while *E. coli* OP50 cultures were grown at 37 °C for two days to compensate for slower growth and lower density. Cultures were washed with PBS and adjusted to OD_600_ = 5.0. Fifty microliters of each culture were spotted on opposite sides of pre-dried 10 cm NGM plates, equidistant from the center, and allowed to dry. For worm preparation, synchronized N2 L1 larvae were plated onto JUb19 or OP50 lawns and grown for 72 h at 20 °C. At 72 h, worms were collected in M9 buffer, and approximately 300 animals per pre-exposure condition were deposited equidistant between the two bacterial spots. Plates were incubated at 20 °C, and worms on each spot were counted every hour for 3 h. A bacterial choice index was calculated as (number on JUb19 − number on OP50) / total number on both spots, where positive values indicate preference for JUb19 and negative values indicate preference for OP50^65^. Data were pooled from eight biological replicates per three experimental replicates per condition, visualized as box-and-whisker plots, and statistical analyses were performed as indicated in the figure legends.

## Acknowledgements

We thank Dr. Emily Troemel, Dr. Damien O’Halloran, Nicole Rangoussis, Paaramitha Warushavithana, and Dustin Howard for valuable comments on the manuscript. We also thank Dr. Michael Herman and Dr. Emily Troemel for providing bacterial and/or *C. elegans* strains. Some strains were provided by the CGC, which is funded by NIH Office of Research Infrastructure Programs (P40 OD010440). The models in Fig. 6 were created using BioRender.com.

## Figures and Figure Legends

**Figure S1:**
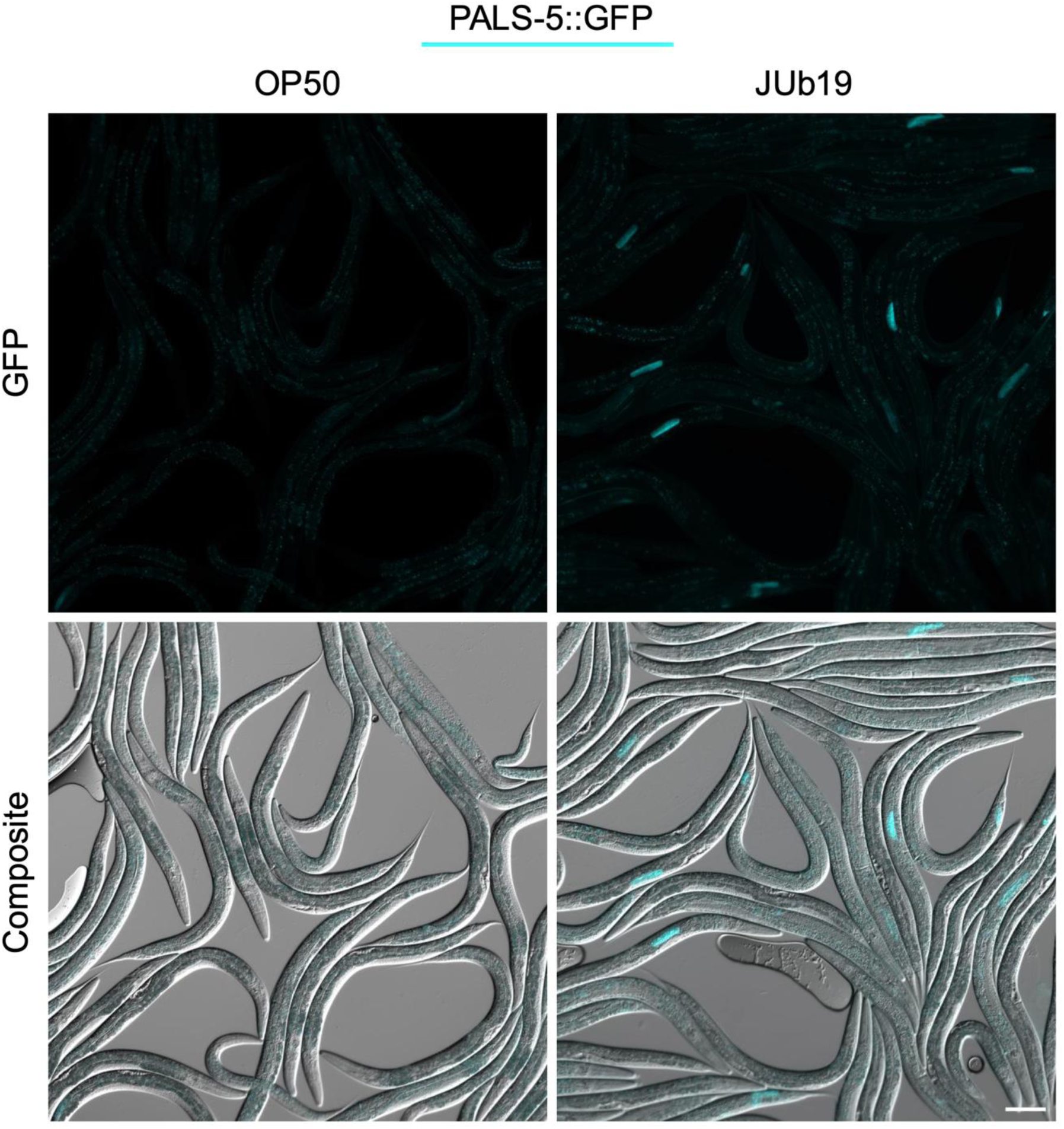
JUb19 induced PALS-5::GFP translational reporter in restricted pattern. Representative images of animals carrying the PALS-5 reporter fed OP50 and JUb19 at 48 h. Composite images consist of merged green channel (PALS-5::GFP) shown in cyan and DIC images. Scale bar: 100 µm

**Figure S2:**
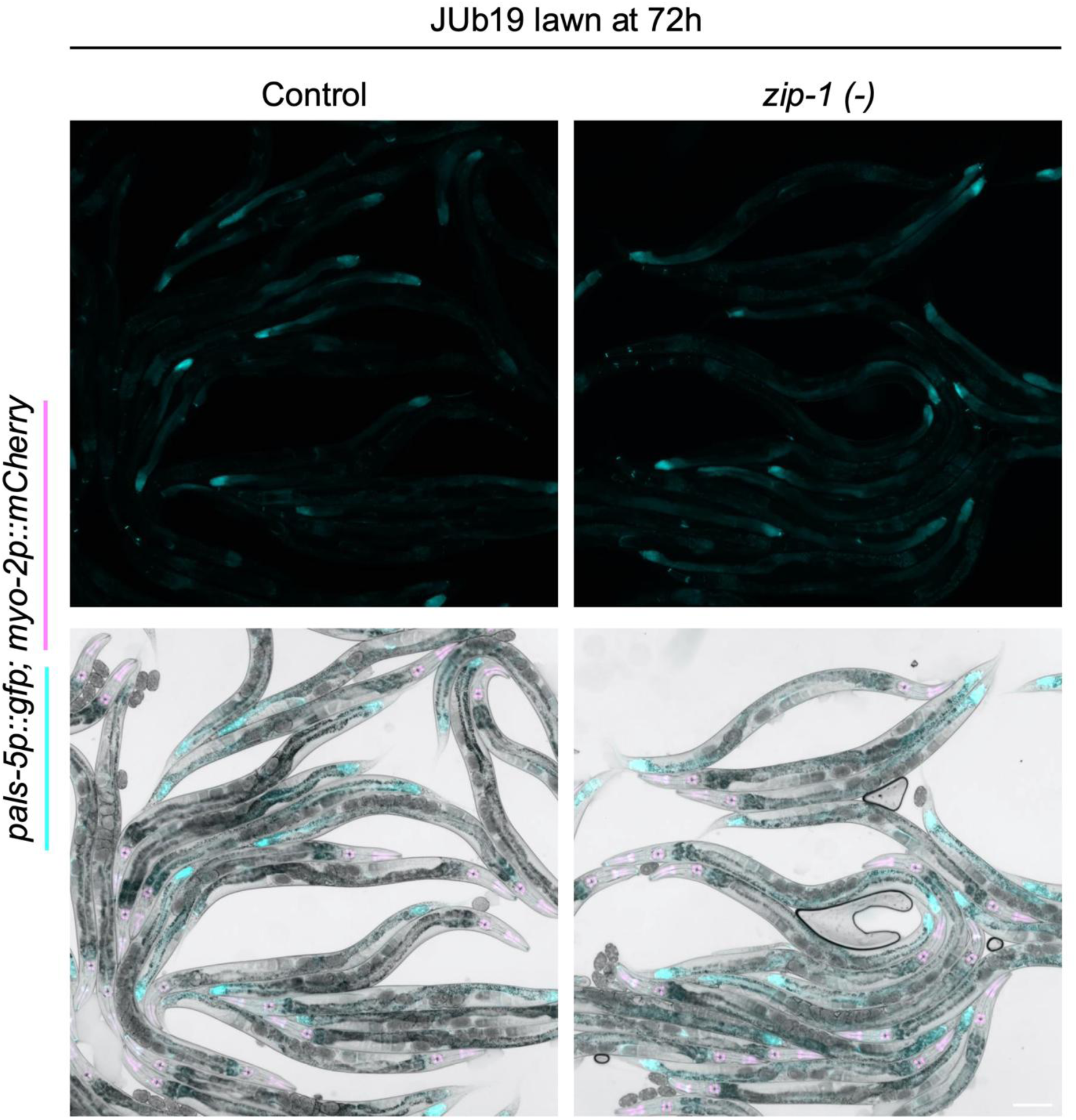
JUb19 IPR reporter induction is not dependent on ZIP-1 following 24h of exposure. Representative images of control and zip-1 mutants with the jyIs8 transgene exposed to JUb19 for 24 h and imaged at 72 h. Composite images consist of merged green channel (*pals-5*p::GFP) shown in cyan, red channel (*myo-2*p::mCherry) shown in magenta, and bright field images. Scale bars: 100 µm

**Figure S3:**
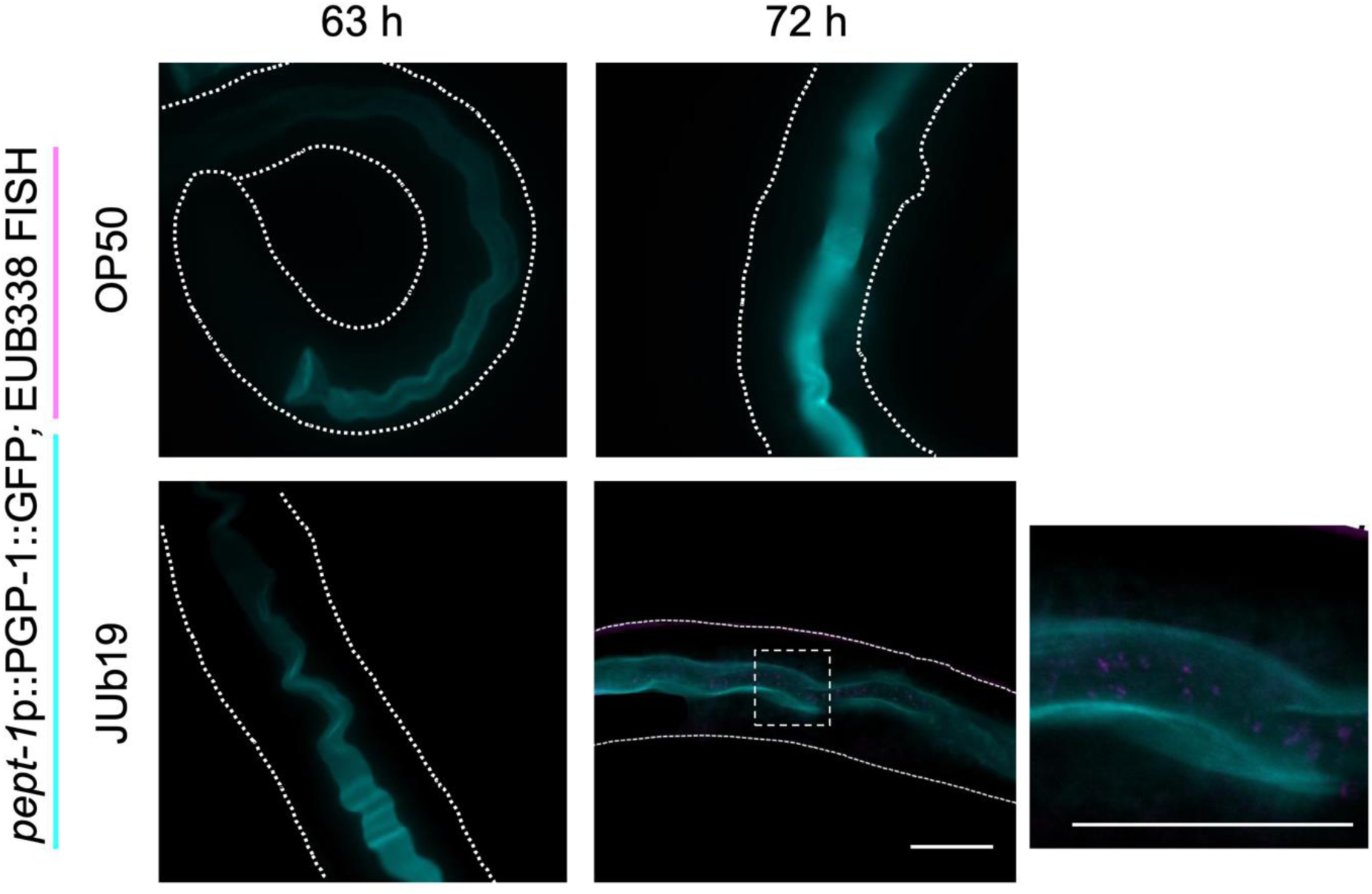
Intestinal colonization at 63 and 72 h. Composite images of *pept-1*p::PGP-1::GFP intestinal apical membrane marker (cyan) and JUb19 labeled with EUB338 red fluorescent probe (magenta). Dotted white lines indicate the outline of each animal. The boxed region in the 72 h JUb19-fed animal is shown enlarged at right to highlight EUB338-stained JUb19 within the intestinal lumen. Scale bar = 100 μm

**Figure S4:**
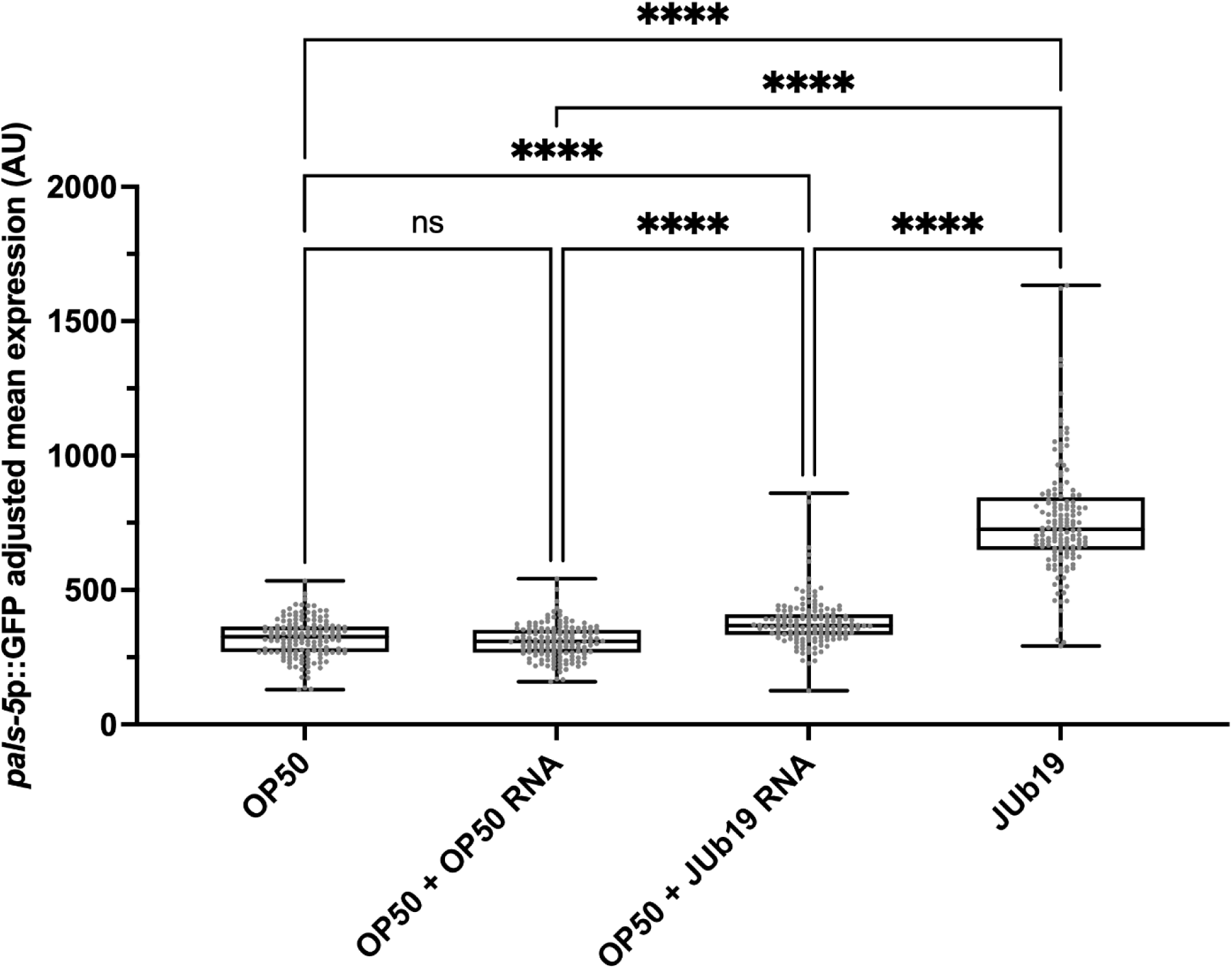
IPR reporter expression analysis following two JUb19 RNA supplementations. JUb19 RNA supplementation assay – reporter expression quantification normalized to background fluorescence. In box-and-whisker plots, the central line marks the median, the box spans the interquartile range (25th–75th percentiles), and the whiskers extend to the smallest and largest values. Gray dots represent measurements for individual animals. Statistical significance was determined using Kruskal-Wallis test for multiple comparisons. *p*-values: OP50 vs OP50+OP50 RNA *p*>0.9999, all remaining comparisons are *p*<0.0001.

**Figure S5:**
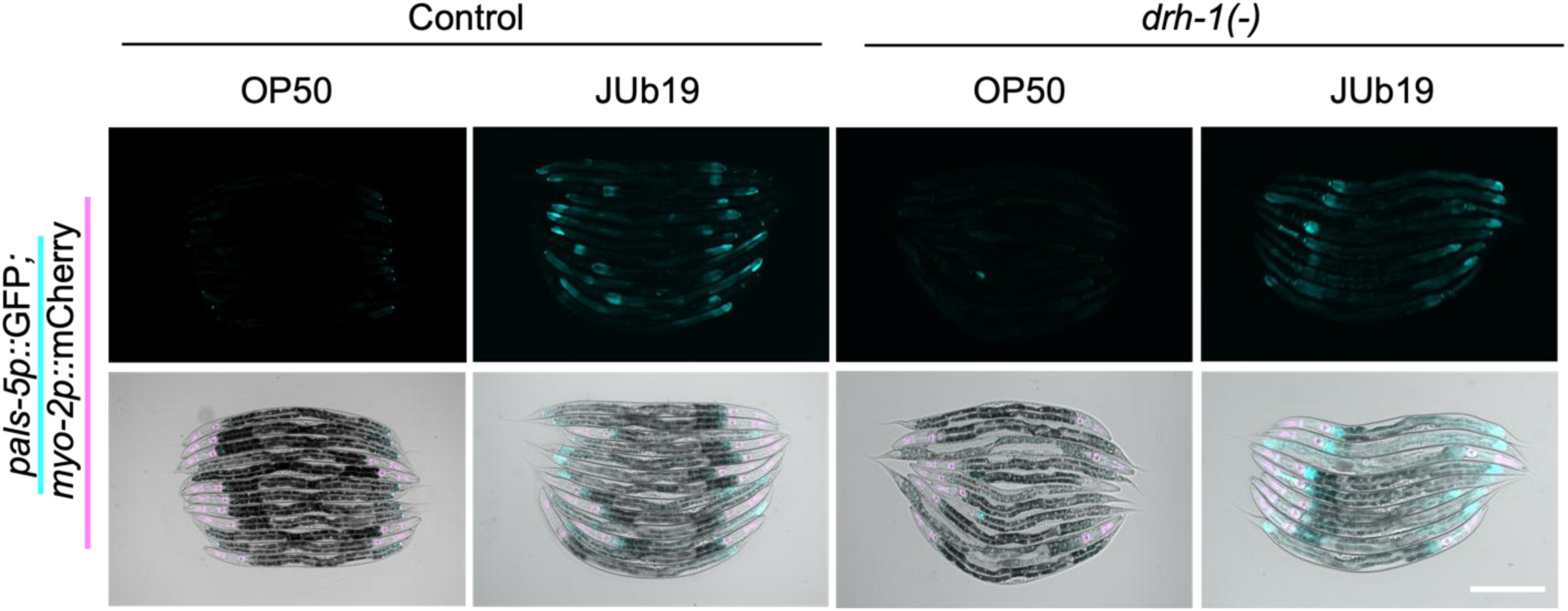
JUb19-RNA induced IPR expression is not dependent on *drh-1*. **a** Representative images of IPR reporter induction in control, and *drh-1* mutants following JUb19 exposure. Composite images consist of merged green channel (*pals-5*p::GFP) shown in cyan, red channel (*myo-2*p::mCherry) shown in magenta, and DIC images. Scale bars: 100 µm.

**Figure S6:**
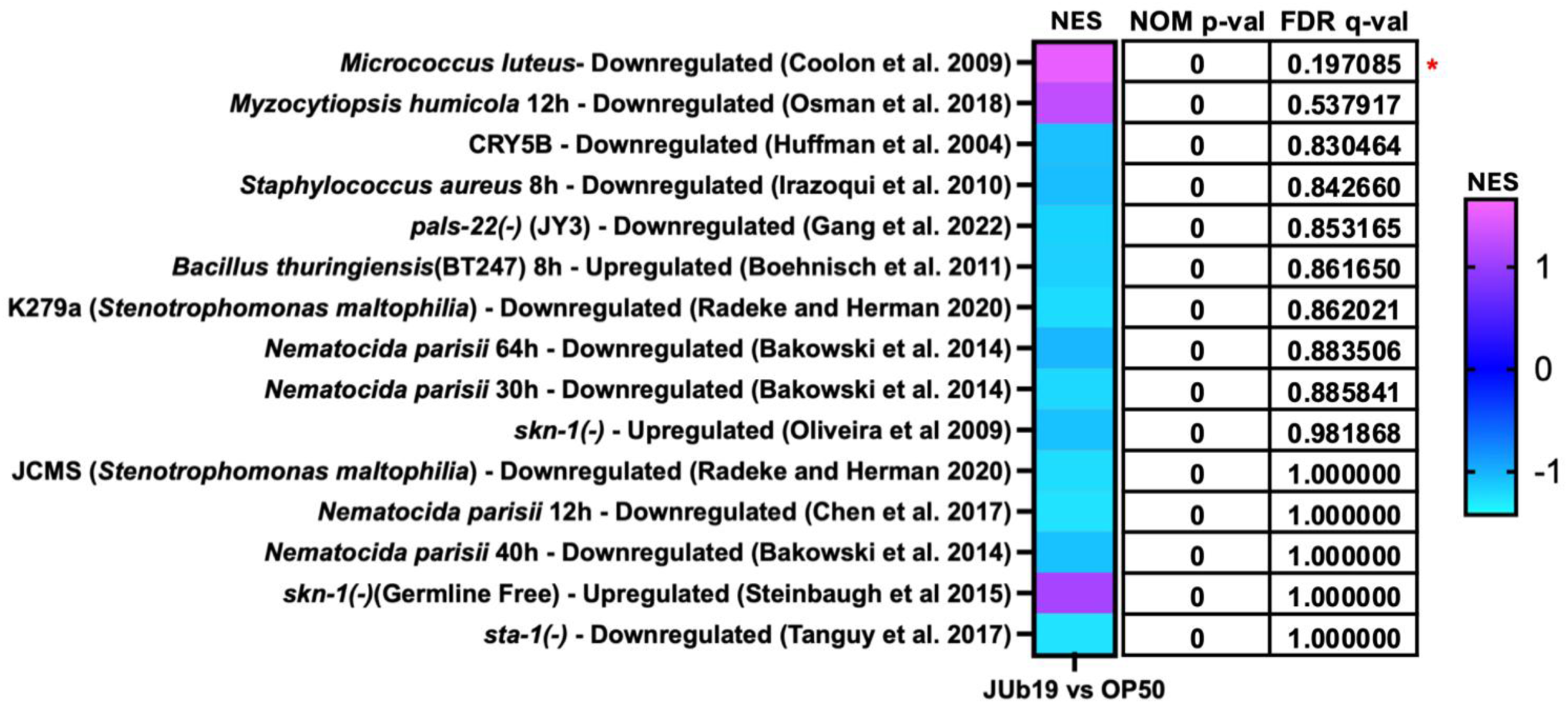
Heat map of GSEA of JUb19 differentially expressed genes. GSEA heat map of immune and stress-related gene sets in JUb19-fed animals compared with OP50 controls. Color indicates normalized enrichment score (NES), with positive values indicating gene sets enriched among genes upregulated in JUb19-fed animals and negative values indicating gene sets enriched among genes downregulated in JUb19-fed animals.

**Figure S7:**
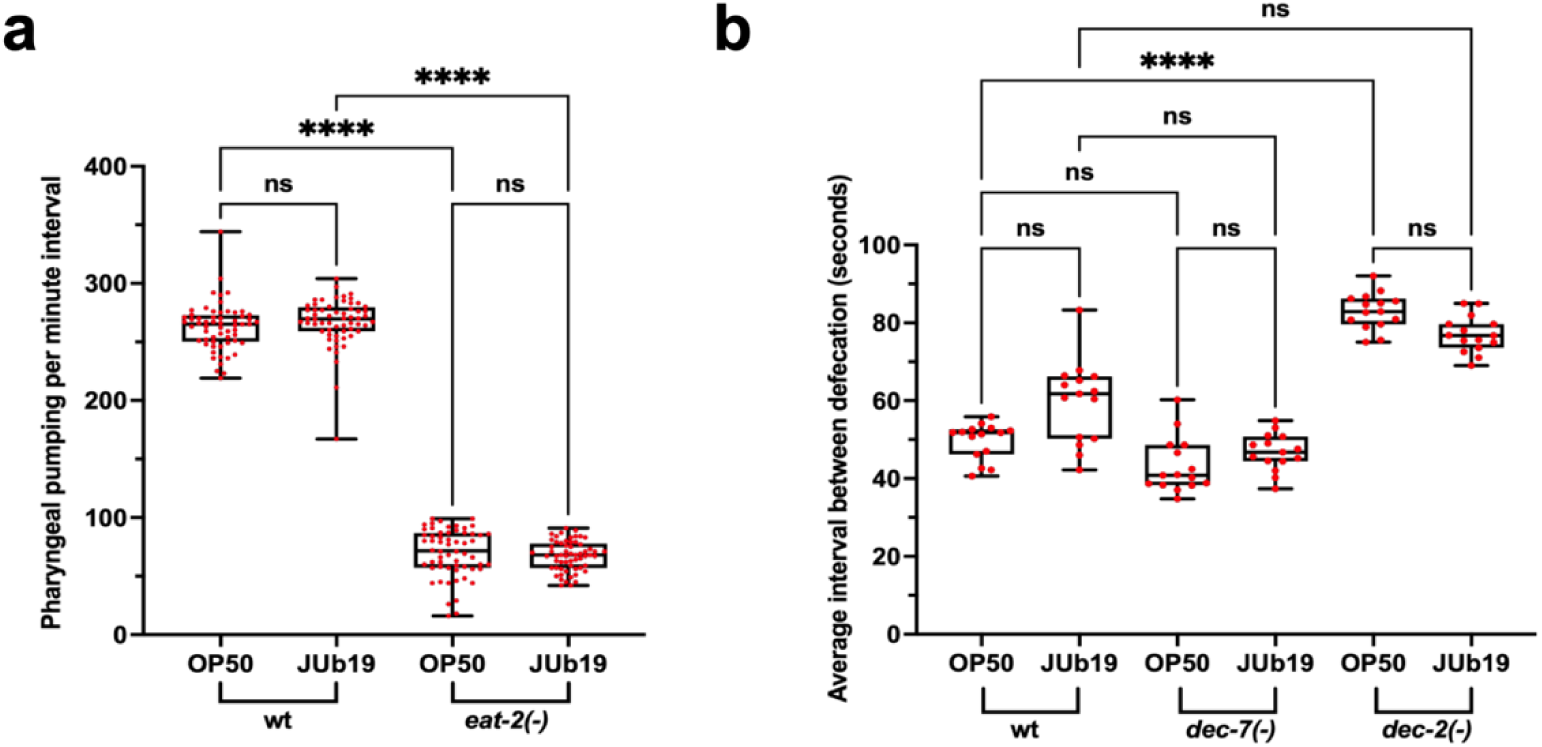
Feeding and defecation rates of JUb19-fed worms. **a** Pharyngeal pumping rates of wild-type and *eat-2(-)* animals on OP50 and JUb19. Pumping was measured in L4-stage worms across three replicates (40 animals/condition/replicate). **b** Defecation rates of wild-type animals and defecation mutants (*dec-7(-)* – faster defecation cycle, and *dec-2(-)* – slower defecation cycle) on OP50 and JUb19. Anal expulsions were measured five gravid adults per condition per three experimental replicates in gravid adults. The average of five defecation cycles was recorded per adult. **a, b** In box-and-whisker plots, the central line marks the median, the box spans the interquartile range (25th–75th percentiles), and the whiskers extend to the smallest and largest values. Red dots represent measurements for individual animals. Statistical significance was determined using the Kruskal-Wallis test for multiple comparisons. *p*-values in (**a**): **** *p*<0.0001, ns non-significant *p*<0.9999. *p*-values in (**b**) are all non-significant *p*<0.9999.

**Supplementary Table 1: Source data.**

**Supplementary Table 2: Differentially expressed genes from RNA seq analysis.**

**Supplementary Table 3: RNA seq normalized counts.**

**Supplementary Table 4: WormCat analysis of RNA seq data.**

**Supplementary Table 5: Comparisons of upregulated genes between JUb19 and three *Stenotrophomonas maltophilia* strains.**

**Supplementary Table 6: GSEA results.**

**Supplementary Table 7: *C. elegans* strains used in this study**.

**Supplementary Table 8: qRT-PCR primers used in this study**.

